# Space is the Place: Effects of Continuous Spatial Structure on Analysis of Population Genetic Data

**DOI:** 10.1101/659235

**Authors:** C.J. Battey, Peter L. Ralph, Andrew D. Kern

**Author notes:** 301 Pacific Hall, University of Oregon Dept. Biology, Institute for Ecology and Evolution. these authors co-supervised this project (Peter L. Ralph, Andrew D. Kern).

## Abstract

Real geography is continuous, but standard models in population genetics are based on discrete, well-mixed populations. As a result many methods of analyzing genetic data assume that samples are a random draw from a well-mixed population, but are applied to clustered samples from populations that are structured clinally over space. Here we use simulations of populations living in continuous geography to study the impacts of dispersal and sampling strategy on population genetic summary statistics, demographic inference, and genome-wide association studies. We find that most common summary statistics have distributions that differ substantially from that seen in well-mixed populations, especially when Wright’s neighborhood size is less than 100 and sampling is spatially clustered. Stepping-stone models reproduce some of these effects, but discretizing the landscape introduces artifacts which in some cases are exacerbated at higher resolutions. The combination of low dispersal and clustered sampling causes demographic inference from the site frequency spectrum to infer more turbulent demographic histories, but averaged results across multiple simulations were surprisingly robust to isolation by distance. We also show that the combination of spatially autocorrelated environments and limited dispersal causes genome-wide association studies to identify spurious signals of genetic association with purely environmentally determined phenotypes, and that this bias is only partially corrected by regressing out principal components of ancestry. Last, we discuss the relevance of our simulation results for inference from genetic variation in real organisms.

## Introduction

The inescapable reality that biological organisms live, move, and reproduce in continuous geography is usually omitted from population genetic models. However, mates tend to live near to one another and to their offspring, leading to a positive correlation between genetic differentiation and geographic distance. This pattern of “isolation by distance” (Wright 1943) is one of the most widely replicated empirical findings in population genetics (Aguillon *et al.* 2017; Jay *et al.* 2012; Sharbel *et al.* 2000). Despite a long history of analytical work describing the genetics of populations distributed across continuous geography (e.g., Wright (1943); Rousset (1997); Barton *et al.* (2002, 2010); Ringbauer *et al.* (2017); Robledo-Arnuncio and Rousset (2010); Wilkins and Wakeley (2002); Wilkins (2004)), much modern work still describes geographic structure as a set of discrete populations connected by migration (e.g., Wright 1931; Epperson 2003; Rousset and Leblois 2011; Shirk and Cushman 2014; Lundgren and Ralph 2019) or as an average over such discrete models (Petkova *et al.* 2015; Al-Asadi *et al.* 2019). For this reason, most population genetics statistics are interpreted with reference to discrete, well-mixed populations, and most empirical papers analyze variation within clusters of genetic variation inferred by programs like *STRUCTURE* (Pritchard *et al.* 2000) with methods that assume these are randomly mating units.

The assumption that populations are “well-mixed” has important implications for downstream inference of selection and demography. Methods based on the coalescent (Kingman 1982; Wakeley 2009) assume that the sampled individuals are a random draw from a well-mixed population that is much larger than the sample (Wakeley and Takahashi 2003). The key assumption is that the individuals of each generation are *exchangeable*, so that there is no correlation between the fate or fecundity of a parent and that of their offspring (Huillet and Möhle 2011). If dispersal or mate selection is limited by geographic proximity, this assumption can be violated in many ways. For instance, if mean viability or fecundity is spatially autocorrelated, then limited geographic dispersal will lead to parent–offspring correlations. Furthermore, nearby individuals will be more closely related than an average random pair, so drawing multiple samples from the same area of the landscape will represent a biased sample of the genetic variation present in the whole population (Städler *et al.* 2009).

Two areas in which spatial structure may be particularly important are demographic inference and genome-wide association studies (GWAS). Previous work has found that discrete population structure can create false signatures of population bottlenecks when attempting to infer demographic histories from microsatellite variation (Chikhi *et al.* 2010), statistics summarizing the site frequency spectrum (SFS) (Ptak and Przeworski 2002; Städler *et al.* 2009; St. Onge *et al.* 2012), or runs of homozygosity in a single individual (Mazet *et al.* 2015). The increasing availability of whole-genome data has led to the development of many methods that attempt to infer detailed trajectories of population sizes through time based on a variety of summaries of genetic data (Liu and Fu 2015; Schiffels and Durbin 2014; Sheehan *et al.* 2013; Terhorst *et al.* 2016). Because all of these methods assume that the populations being modeled are approximately randomly mating, they are likely affected by spatial biases in the genealogy of sampled individuals (Wakeley 1999), which may lead to incorrect inference of population changes over time (Mazet *et al.* 2015). However, previous investigations of these effects have focused on discrete rather than continuous space models, and the level of isolation by distance at which inference of population size trajectories become biased by structure is not well known. Here we test how two methods suitable for use with large samples of individuals – stairwayplot (Liu and Fu 2015) and SMC++ (Terhorst *et al.* 2016) – perform when applied to populations evolving in continuous space with varying sampling strategies and levels of dispersal.

Spatial structure is also a major challenge for interpreting the results of genome-wide association studies (GWAS). This is because many phenotypes of interest have strong geographic differences due to the (nongenetic) influence of environmental or socioeconomic factors, which can therefore show spurious correlations with spatially patterned allele frequencies (Bulik-Sullivan *et al.* 2015; Mathieson and McVean 2012). Indeed, two recent studies found that previous evidence of polygenic selection on human height in Europe was confounded by subtle population structure (Sohail *et al.* 2018; Berg *et al.* 2018), suggesting that existing methods to correct for population structure in GWAS are insufficient. However we have little quantitative idea of the population and environmental parameters that can be expected to lead to biases in GWAS.

Last, some of the most basic tools of population genetics are summary statistics like *F*_*IS*_ and Tajima’s *D*, which are often interpreted as reflecting the influence of selection or demography on sampled populations (Tajima 1989). Statistics like Tajima’s *D* are essentially summaries of the site frequency spectrum, which itself reflects variation in branch lengths and tree structure of the underlying genealogies of sampled individuals. Geographically limited mate choice distorts the distribution of these genealogies (Maruyama 1972; Wakeley 1999), which can affect the value of Tajima’s *D* (Städler *et al.* 2009). Similarly, the distribution of tract lengths of identity by state among individuals contains information about not only historical demography (Harris and Nielsen 2013; Ralph and Coop 2013) and selection (Garud *et al.* 2015), but also dispersal and mate choice (Ringbauer *et al.* 2017; Baharian *et al.* 2016). We are particularly keen to examine how such summaries will be affected by models that incorporate continuous space, both to evaluate the assumptions underlying existing methods and to identify where the most promising signals of geography lie.

To study this, we have implemented an individual-based model in continuous geography that incorporates overlapping generations, local dispersal of offspring, and density-dependent survival. We simulate chromosome-scale genomic data in tens of thousands of individuals from parameter regimes relevant to common subjects of population genetic investigation such as humans and *Drosophila*, and output the full genealogy and recombination history of all final-generation individuals. We use these simulations to test how sampling strategy interacts with geographic population structure to cause systematic variation in population genetic summary statistics typically analyzed assuming discrete population models. We then examine how the fine-scale spatial structures occurring under limited dispersal impact demographic inference from the site frequency spectrum. Last, we examine the impacts of continuous geography on genome-wide association studies (GWAS) and identify regions of parameter space under which the results from GWAS may be misleading.

## Materials and Methods

### Modeling Evolution in Continuous Space

The degree to which genetic relationships are geographically correlated depends on the chance that two geographically nearby individuals are close relatives – in modern terms, by the tension between migration (the chance that one is descended from a distant location) and coalescence (the chance that they share a parent). A key early observation by Wright (1946) is that this balance is often nicely summarized by the “neighborhood size”, defined to be *N*_*W*_ = 4*πρσ*^2^, where *σ* is the mean parent– offspring distance and *ρ* is population density. This can be thought of as proportional to the average number of potential mates for an individual (those within distance 2*σ*), or the number of potential parents of a randomly chosen individual. Empirical estimates of neighborhood size vary hugely across species – even in human populations, estimates range from 40 to over 5,000 depending on the population and method of estimation (Table 1).

**Table 1.** Neighborhood size estimates from empirical studies.

| Species | Description | Neighborhood Size | Method | Citation |
| --- | --- | --- | --- | --- |
| *Ipomopsis aggregata* | flowering plant | 12.60 - 37.80 | Genetic | (Campbell and Dooley 1992) |
| *Borrchia frutescens* | salt marsh plant | 20 - 30 | Genetic+Survey | (Antlfinger 1982) |
| *Oreamnos americanus* | mountain goat | 36 - 100 | Genetic | (Shirk and Cushman 2014) |
| *Homo sapiens* | Gainj- and Kalam-speaking people, Papua New Guinea | 40 - 213 | Genetic | (Rousset 1997) |
| *Formica sp.* | colonial ants | 50 - 100 | Genetic | (Pamilo 1983) |
| *Astrocaryum mexicanum* | palm tree | 102 - 895 | Genetic+survey | (Eguarte *et al.* 1993) |
| *Spermophilus mollis* | ground squirrel | 204 - 480 | Genetic+Survey | (Antolin *et al.* 2001) |
| *Sceloporus olivaceus* | lizard | 225 - 270 | Survey | (Kerster 1964) |
| *Dieffenbachia longispatha* | beetle-pollinated colonial herb | 227 - 611 | Survey | (Young 1988) |
| *Aedes aegypti* | Yellow-fever mosquito | 268 | Genetic | (Jasper *et al.* 2019) |
| *Homo sapiens* | Gainj- and Kalam-speaking people, Papua New Guinea | 410 | Survey | (Rousset 1997) |
| *Quercus laevis* | Oak tree | > 440 | Genetic | (Berg and Hamrick 1995) |
| *Drosophila pseudoobscura* | fruit fly | 500 - 1,000 | Survey+Crosses | (Wright 1946) |
| *Homo sapiens* | POPRES data NE Europe | 1,342 - 5,425 | Genetic | (Ringbauer *et al.* 2017) |
| *Bebicium vittatum* | intertidal snail | 240,000 | Survey | (Rousset 1997) |
| *Bebicium vittatum* | intertidal snail | 360,000 | Genetic | (Rousset 1997) |

The first approach to modeling continuously distributed populations was to endow individuals in a Wright-Fisher model with locations in continuous space. However, since the total size of the population is constrained, this introduces interactions between arbitrarily distant individuals, which (aside from being implausible) was shown by Felsenstein (1975) to eventually lead to unrealistic population clumping if the range is sufficiently large. Another method for modeling spatial populations is to assume the existence of a grid of discrete randomly mating populations connected by migration, thus enforcing regular population density by edict. Among many other results drawn from this class of “lattice” or “stepping stone” models (Epperson 2003), Rousset (1997) showed that the slope of the linear regression of genetic differentiation (*F*_*ST*_) against the logarithm of spatial distance is an estimate of neighborhood size. Although these grid models may be good approximations of continuous geography in many situations, they do not model demographic fluctuations, and limit investigation of spatial structure below the level of the deme, assumptions whose impacts are unknown. An alternative method for dealing with continuous geography is a new class of coalescent models, the Spatial Lambda Fleming-Viot models (Barton *et al.* 2010; Kelleher *et al.* 2014).

To avoid hard-to-evaluate approximations, we here used forward-time, individual-based simulations across continuous geographical space. The question of what regulates real populations has a long history and many answers (e.g., Lloyd 1967; Antonovics and Levin 1980; Crawley 1990), but it is clear that populations must at some point have density-dependent feedback on population size, or else they would face eventual extinction or explosion. In the absence of unrealistic global population regulation, this regulation must be local, and there are many ways to achieve this (Bolker *et al.* 2003). In our simulations, each individual’s probability of survival is a decreasing function of local population density, which shifts reproductive output towards low-density regions, and produces total census sizes that fluctuate around an equilibrium. This also prevents the population clumping seen by Felsenstein (1975) (Supplemental Figure S1)). Such models have been used extensively in ecological modeling (Durrett and Levin 1994; Bolker and Pacala 1997; Law *et al.* 2003; Fournier and Méléard 2004; Champer *et al.* 2019) but rarely in population genetics, where to our knowledge implementations of continuous space models before their availability through SLiM (Haller and Messer 2019) have focused on a small number of genetic loci (e.g., Slatkin and Barton 1989; Barton *et al.* 2002; Robledo-Arnuncio and Rousset 2010; Rossine 2014; Jackson and Fahrig 2014), which limits the ability to investigate the impacts of continuous space on genome-wide genetic variation as is now routinely sampled from real organisms. By simulating chromosome-scale sequence alignments and complete population histories we are able to treat our simulations as real populations and replicate the sampling designs and analyses commonly conducted on real genomic data.

### A Forward-Time Model of Evolution in Continuous Space

We simulated populations using the program SLiM v3.1 (Haller and Messer 2019). Each time step consists of three stages: reproduction, dispersal, and mortality. To reduce the parameter space we use the same parameter, denoted *σ*, to modulate the spatial scale of interactions at all three stages by adjusting the standard deviation of the corresponding Gaussian functions. As in previous work (Wright 1943; Ringbauer *et al.* 2017), *σ* is equal to the mean parent-offspring distance.

At the beginning of the simulation individuals are distributed uniformly at random on a continuous, square landscape. Individuals are hermaphroditic, and each time step, each produces a Poisson number of offspring with mean 1/*L*. Offspring disperse a Gaussian-distributed distance away from the parent with mean zero and standard deviation *σ* in both the *x* and *y* coordinates. Each offspring is produced with a mate selected randomly from those within distance 3*σ*, with probability of choosing a neighbor at distance *d* proportional to exp(−*d*^2^/2*σ*^2^).

To maintain a stable population, mortality increases with local population density. To do this we say that individuals at distance *d* have a competitive interaction with strength *g*(*d*), where *g* is the Gaussian density with mean zero and standard deviation *σ*. Then, the sum of all competitive interactions with individual *i* is *n*_*i*_ = ∑_*j*_ *g*(*d*_*ij*_), where *d*_*ij*_ is the distance between individuals *i* and *j* and the sum is over all neighbors within distance 3*σ*. Since *g* is a probability density, *n*_*i*_ is an estimate of the number of nearby individuals per unit area. Then, given a per-unit carrying capacity *K*, the probability of survival until the next time step for individual *i* is

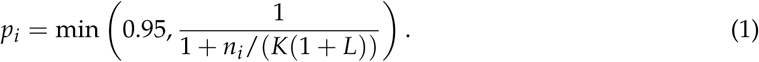

We chose this functional form so that the equilibrium population density per unit area is close to *K*, and the mean lifetime is around *L*; for more description see the Appendix.

An important step in creating any spatial model is dealing with range edges. Because local population density is used to model competition, edge or corner populations can be assigned artificially high fitness values because they lack neighbors within their interaction radius but outside the bounds of the simulation. We approximate a decline in habitat suitability near edges by decreasing the probability of survival proportional to the square root of distance to edges in units of *σ*. The final probability of survival for individual *i* is then

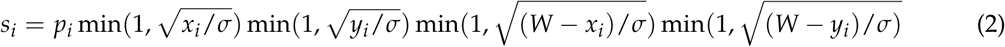

where *x*_*i*_ and *y*_*i*_ are the spatial coordinates of individual *i*, and *W* is the width (and height) of the square habitat. This buffer roughly counteracts the increase in fitness individuals close to the edge would otherwise have, though the effect is relatively subtle (Figure S2).

To isolate spatial effects from other components of the model such as overlapping generations, increased variance in reproductive success, and density-dependent fitness, we also implemented simulations identical to those above except that mates are selected uniformly at random from the population, and offspring disperse to a uniform random location on the landscape. We refer to this model as the “random mating” model, in contrast to the first, “spatial” model.

We stored the full genealogy and recombination history of final-generation individuals as tree sequences (Kelleher *et al.* 2018), as implemented in SLiM (Haller *et al.* 2019). Scripts for figures and analyses are available at https://github.com/kern-lab/spaceness.

We ran 400 simulations for the spatial and random-mating models on a square landscape of width *W* = 50 with per-unit carrying capacity *K* = 5 (census *N* ≈ 10, 000), average lifetime *L* = 4, genome size 10^8^bp, recombination rate 10^−9^ per bp per generation, and drawing *σ* values from a uniform distribution between 0.2 and 4. To speed up the simulations and limit memory overhead we used a mutation rate of 0 in SLiM and later applied mutations to the tree sequence with msprime’s `mutate` function (Kelleher *et al.* 2016). Because msprime applies mutations proportionally to elapsed time, we divided the mutation rate of 10^−8^ mutations per site per generation by the average generation time estimated for each value of *σ* (see ‘Demographic Parameters’ below) to convert the rate to units of mutations per site per unit time. We verify that this procedure produces the same site frequency spectrum as applying mutations directly in SLiM in Figure S3, in agreement with theory (Ralph *et al.* 2019). Simulations were run for 1.6 million timesteps (approximately 30*N* generations).

We also compared our model’s output to a commonly-used approximation of continuous space, the stepping-stone model, which we simulated with `msprime` (Kelleher *et al.* 2016). These results are discussed in detail in the Appendix, but in general we find that the demographic structure of a stepping-stone model can depend strongly on the chosen discretization, and some artifacts of discretization seem to become stronger in the limit of a fine grid. For many summary statistics, finer discretizations (we used a 50×50 grid) produced similar results to the continuous model, but this was not true for others (e.g., *F*_*IS*_ and Tajima’s *D*), which differed from the continuous model *more* at finer discretizations.

### Demographic Parameters

Our demographic model includes parameters that control population density (*K*), mean life span (*L*), and dispersal distance (*σ*). However, nonlinearity of local demographic stochasticity causes actual realized averages of these demographic quantitites to deviate from the specified values in a way that depends on the neighborhood size. Therefore, to properly compare to theoretical expectations, we empirically calculated these demographic quantities in simulations. We recorded the census population size in all simulations, and used mean population density (*ρ*, census size divided by total area) to compute neighborhood size as *N*_*W*_ = 4*πρσ*^2^. To estimate generation times, we stored ages of the parents of every new individual born across 200 timesteps, after a 100 generation burn-in, and took the mean. To estimate variance in offspring number, we tracked the lifetime total number of offspring for all individuals for 100 timesteps following a 100-timestep burn-in period, and calculated the variance in number of offspring across all individuals in timesteps 50-100. All calculations were performed with information recorded in the tree sequence, using pyslim (Peter L Ralph and Ashander ????).

### Sampling

Our model records the genealogy and sequence variation of the complete population, but in real data, genotypes are only observed from a relatively small number of sampled individuals. We modeled three sampling strategies similar to common data collection methods in empirical genetic studies (Figure 1). “Random” sampling selects individuals at random from across the full landscape, “point” sampling selects individuals proportional to their distance from four equally spaced points on the landscape, and “midpoint” sampling selects individuals in proportion to their distance from the middle of the landscape. Downstream analyses were repeated across all sampling strategies.

**Figure 1.**
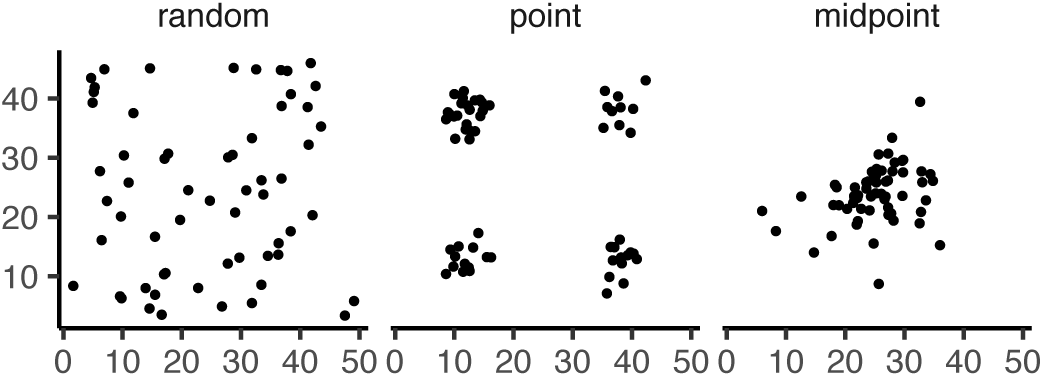
Example sampling maps for 60 individuals on a 50×50 landscape for midpoint, point, and random sampling strategies, respectively.

### Summary Statistics

We calculated the site frequency spectrum and a set of 18 summary statistics (Table S1) from 60 diploid individuals sampled from the final generation of each simulation using the python package scikit-allel (Miles and Harding 2017). Statistics included common single-population summaries including mean pairwise divergence (*π*), inbreeding coefficient (*F*_*IS*_), and Tajima’s *D*, as well as (motivated by Rousset (1997)’s results) the correlation coefficient between the logarithm of the spatial distance and the proportion of identical base pairs across pairs of individuals.

Following recent studies that showed strong signals for dispersal and demography in the distribution of shared haplotype block lengths (e.g., Ringbauer *et al.* 2017; Baharian *et al.* 2016), we also calculated various summaries of the distribution of pairwise identical-by-state (IBS) block lengths among sampled chromosomes, defined to be the set of distances between adjacent sites that differ between the two chromosomes. The full distribution of lengths of IBS tracts for each pair of chromosomes was first calculated with a custom python function. We then calculated the first three moments of this distribution (mean, variance, and skew) and the number of blocks over 10^6^ base pairs both for each pair of individuals and for the full distribution across all pairwise comparisons. We then calculated correlation coefficients between spatial distance and each moment of the pairwise IBS tract distribution. Because more closely related individuals on average share longer haplotype blocks we expect that spatial distance will be negatively correlated with mean haplotype block length, and that this correlation will be strongest (i.e., most negative) when dispersal is low. The variance, skew, and count of long haplotype block statistics are meant to reflect the relative length of the right (upper) tail of the distribution, which represents the frequency of long haplotype blocks, and so should reflect recent demographic events (Chapman and Thompson 2002). For a subset of simulations, we also calculated cumulative distributions for IBS tract lengths across pairs of distant (more than 40 map units) and nearby (less than 10 map units) individuals. Last, we examined the relationship between allele frequency and the spatial dispersion of an allele by calculating the average distance between individuals carrying each derived allele.

The effects of sampling on summary statistic estimates were summarized by testing for differences in mean (ANOVA, (R Core Team 2018)) and variance (Levene’s test, (Fox and Weisberg 2011)) across sampling strategies for each summary statistic.

### Demographic Inference

To assess the impacts of continuous spatial structure on demographic inference we inferred population size histories for all simulations using two approaches: stairwayplot (Liu and Fu 2015) and SMC++ (Terhorst *et al.* 2016). Stairwayplot fits its model to a genome-wide estimate of the SFS, while SMC++ also incorporates linkage information. For both methods we sampled 20 individuals from all spatial simulations using random, midpoint, and point sampling strategies.

As recommended by its documentation, we used stairwayplot to fit models with multiple bootstrap replicates drawn from empirical genomic data, and took the median inferred *N*_*e*_ per unit time as the best estimate. We calculated site frequency spectra with scikit-allel (Miles and Harding 2017), generated 100 bootstrap replicates per simulation by resampling over sites, and fit models for all bootstrap samples using default settings.

For SMC++, we first output genotypes as VCF with msprime and then used SMC++’s standard pipeline for preparing input files assuming no polarization error in the SFS. We used the first individual in the VCF as the “designated individual” when fitting models, and allowed the program to estimate the recombination rate during optimization. We fit models using the ‘estimate’ command rather than the now recommended cross-validation approach because our simulations had only a single contig.

To evaluate the performance of these methods we binned simulations by neighborhood size, took a rolling median of inferred *N*_*e*_ trajectories across all model fits in a bin for each method and sampling strategy. We also examined how varying levels of isolation by distance impacted the variance of *N*_*e*_ estimates by calculating the standard deviation of *N*_*e*_ from each best-fit model.

### Association Studies

To assess the degree to which spatial structure confounds GWAS we simulated four types of nongenetic phenotype variation for 1000 randomly sampled individuals in each spatial SLiM simulation and conducted a linear regression GWAS with principal components as covariates in PLINK (Purcell *et al.* 2007). SNPs with a minor allele frequency less than 0.5% were excluded from this analysis. Phenotype values were set to vary by two standard deviations across the landscape in a rough approximation of the variation seen in height across Europe (Turchin *et al.* 2012; Garcia and Quintana-Domeque 2006, 2007). Conceptually our approach is similar to that taken by Mathieson and McVean (2012), though here we model fully continuous spatial variation and compare GWAS output across a range of dispersal distances.

In all simulations, the phenotype of each individual is determined by drawing from a Gaussian distribution with standard deviation 10 and a mean that may depend on spatial position. In spatially varying models, the mean phenotype differs by two standard deviations across the landscape. We then adjust the geographic pattern of mean phenotype to create four types of spatially autocorrelated environmental influences on phenotype. In the first simulation of *nonspatial* environments, the mean did not change, so that all individuals’ phenotypes were drawn independently from a Gaussian distribution with mean 110 and standard deviation 10. Next, to simulate *clinal* environmental influences on phenotype, we increased the mean phenotype from 100 on the left edge of the range to 120 on the right edge (two phenotypic standard deviations). Concretely, the mean phenotype *p* for an individual at position (*x, y*) is *p* = 100 + 2*x*/5. Third, we simulated a more concentrated “*corner*” environmental effect by setting the mean phenotype to 120 for individuals with both *x* and *y* coordinates below 20 (two standard deviations above the rest of the map). Finally, in “*patchy*” simulations we selected 10 random points on the map and set the mean phenotype of all individuals within three map units of each of these points to 120.

We performed principal components analysis (PCA) using scikit-allel (Miles and Harding 2017) on the matrix of derived allele counts by individual for each simulation. SNPs were first filtered to remove strongly linked sites by calculating LD between all pairs of SNPs in a 200-SNP moving window and dropping one of each pair of sites with an *R*^2^ over 0.1. The LD-pruned allele count matrix was then centered and all sites scaled to unit variance when conducting the PCA, following recommendations in Patterson *et al.* (2006).

We ran linear-model GWAS both with and without the first 10 principal components as covariates in PLINK and summarized results across simulations by counting the number of SNPs with *p*-value below 0.05 after adjusting for an expected false positive rate of less than 5% (Benjamini and Yekutieli 2001). We also examined *p* values for systematic inflation by comparing to the values expected from a uniform distribution (because no SNPs were used when generating phenotypes, well-calibrated *p*-values should be uniform).

Results from all analyses were summarized and plotted with the “ggplot2” (Wickham 2016) and “cowplot” (Wilke 2019) packages in R (R Core Team 2018).

## Results

### Demographic Parameters and Run Times

Adjusting the spatial dispersal and interaction distance, *σ*, has a surprisingly large effect on demographic quantities that are usually fixed in Wright-Fisher models – the generation time, census population size, and variance in offspring number, shown in Figure 2. Because our simulation is paramaterized on an individual level, these population parameters emerge as a property of the interactions among individuals rather than being directly set. Variation across runs occurs because, even though the parameters *K* and *L* that control population density and mean lifetime respectively were the same in all simulations, the strength of stochastic effects depends strongly on the spatial interaction distance *σ*. For instance, the population density near to individual *i* (denoted *n*_*i*_ above) is computed by averaging over roughly *N*_*W*_ = 4*πKσ*^2^ individuals, and so has standard deviation proportional to 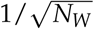 – it is more variable at lower densities. (Recall that *N*_*W*_ is Wright’s neighborhood size.) Since the probability of survival is a nonlinear function of *n*_*i*_, actual equilibrium densities and lifetimes differ from *K* and *L*. This is the reason that we included *random mating* simulations – where mate choice and offspring dispersal are both nonspatial – since this should preserve the random fluctuations in local population density while destroying any spatial genetic structure. We verified that random mating models retained no geographic signal by showing that summary statistics did not differ significantly between sampling regimes (Table S2), unlike in spatial models (discussed below).

**Figure 2.**
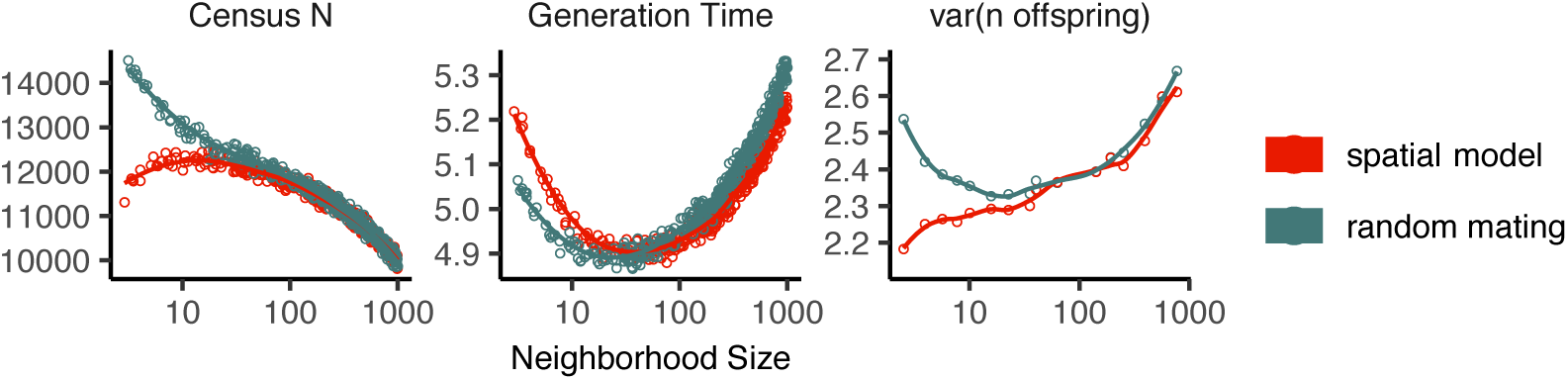
Genealogical parameters from spatial and random mating SLiM simulations, by neighborhood size.

There are a few additional things to note about Figure 2. First, all three quantities are non-monotone with neighborhood size. Census size largely declines as neighborhood size increases for both the spatial and random mating models. However, for spatial models this decline only begins for neighborhood size ≥ 10. Spatial and random mating models are indistinguishable from one another for neighborhood sizes larger than 100. Census sizes range from around 14,000 at low *σ* in the random mating model to 10,000 for both models when neighborhood sizes approach 1,000. The scaling of census sizes in both random-mating and spatial models appears to be related to two consequences of the spatial competition function: the decline of fitness at range edges, which effectively reduces the habitable area by one *σ* around the edge of the map and so results in a smaller habitable area at high *σ* values; and variation in the equilibrium population density given varying competition radii. Furthermore, census size increases in spatial models as neighborhood size increases from 2 to 10. This may reflect an Allee effect (Allee *et al.* 1949) in which some individuals are unable to find mates when the mate selection radius is very small.

Generation time similarly shows complex behavior with respect to neighborhood sizes, and varies between 5.2 and 4.9 timesteps per generation across the parameter range explored. Under both the spatial and random mating models, generation time reaches a minimum at a neighborhood size of around 50. Interestingly, under the range of neighborhood sizes that we examined, generation times between the random mating and spatial models are never quite equivalent – presumably this would cease to be the case at neighborhood sizes higher than we simulated here.

Last, we looked at the variance in number of offspring – a key parameter determining the effective population size. Surprisingly, the spatial and random mating models behave quite differently: while the variance in offspring number increases nearly monotonically under the spatial model, the random mating model actually shows a decline in the variance in offspring number until a neighborhood size of around 10 before it increases and eventually equals what we observe in the spatial case.

Run times for our model scale approximately linearly with neighborhood size (Figure 3), with the lowest neighborhood sizes reaching 30N generations in around a day and those with neighborhood size approaching 1,000 requiring up to three weeks of computation. As currently implemented running simulations at neighborhood sizes more than 1,000 to coalescence is likely impractical, though running these models for more limited timescales and then “recapitating” the simulation using reverse-time simulation from the resulting tree sequence in msprime is possible (Haller *et al.* 2019).

**Figure 3.**
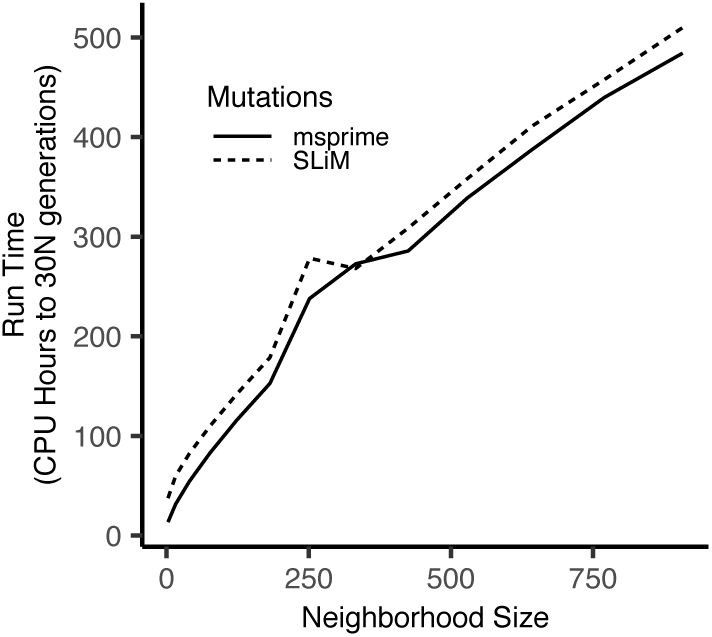
Run times of continuous space simulations with landscape width 50 and expected density 5 under varying neighborhood size. Times are shown for simulations run with mutations applied directly in SLiM (dashed lines) or later applied to tree sequences with msprime (solid lines). Times for simulations run with tree sequence recording disabled are shown in grey.

### Impacts of Continuous Space on Population Genetic Summary Statistics

Even though certain aspects of population demography depend on the scale of spatial interactions, it still could be that population genetic variation is well-described by a well-mixed population model. Indeed, mathematical results suggest that genetic variation in some spatial models should be well-approximated by a Wright-Fisher population if neighborhood size is large and all samples are geographically widely separated (**??**). However, the behavior of most common population genetic summary statistics other than Tajima’s *D* (Städler *et al.* 2009) has not yet been described in realistic geographic models. Moreover, as we will show, spatial sampling strategies can affect summaries of genetic variation at least as strongly as the underlying population dynamics.

#### Site Frequency Spectra and Summaries of Diversity

Figure 4 shows the effect of varying neighborhood size and sampling strategy on the site frequency spectrum (Figure 4, Figure S5) and several standard population genetic summary statistics (Figure 4B; additional statistics are shown in Figure S4). Consistent with findings in island and stepping stone simulations (Städler *et al.* 2009), the SFS shows a significant enrichment of intermediate frequency variants in comparison to the nonspatial expectation. This bias is most pronounced below a neighborhood size of 100 and is exacerbated by midpoint and point sampling of individuals (depicted in Figure 1). Reflecting this, Tajima’s *D* is quite positive in the same situations (Figure 4B). Notably, the point at which Tajima’s *D* approaches 0 differs strongly across sampling strategies – varying from a neighborhood size of roughly 50 for random sampling to at least 1000 for midpoint sampling.

**Figure 4.**
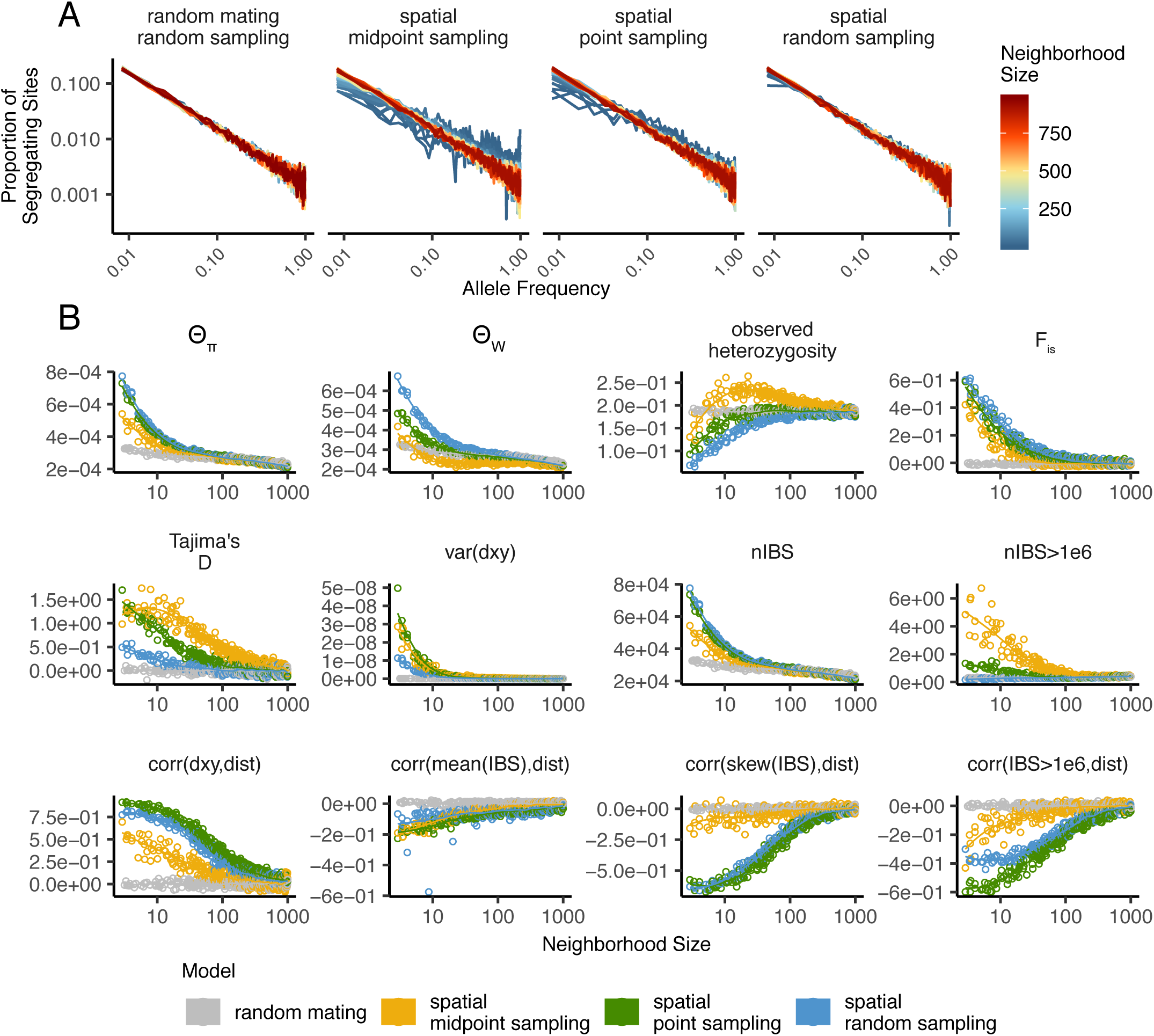
Site frequency spectrum (A; note axes are log-scaled) and summary statistic distributions (B) by sampling strategy and neighborhood size.

One of the most commonly used summaries of variation is Tajima’s summary of nucleotide diversity, *θ*_*π*_, calculated as the mean density of nucleotide differences averaged across pairs of samples. As can be seen in Figure 4B, *θ*_*π*_ in the spatial model is inflated by up to three-fold relative to the random mating model. This pattern is opposite the expectation from census population size (Figure 2), because the spatial model has *lower* census size than the random mating model at neighborhood sizes less than 100. Differences between these models likely occur because *θ*_*π*_ is a measure of mean time to most recent common ancestor between two samples, and at small values of *σ*, the time for dispersal to mix ancestry across the range exceeds the mean coalescent time under random mating. (For instance, at the smallest value of *σ* = 0.2, the range is 250 dispersal distances wide, and since the location of a diffusively moving lineage after *k* generations has variance *Kσ*^2^, it takes around 250^2^ = 62500 generations to mix across the range, which is roughly ten times larger than the random mating effective population size). *θ*_*π*_ using each sampling strategy approaches the random mating expectation at its own rate, but by a neighborhood size of around 100 all models are roughly equivalent. Interestingly, the effect of sampling strategy is reversed relative to that observed in Tajima’s D – midpoint sampling reaches random mating expectations around neighborhood size 50, while random sampling is inflated until around neighborhood size 100.

Values of observed heterozygosity and its derivative *F*_*IS*_ also depend heavily on neighborhood size under spatial models as well as the sampling scheme. *F*_*IS*_ is inflated above the expectation across most of the parameter space examined and across all sampling strategies. This effect is caused by a deficit of heterozygous individuals in low-dispersal simulations – a continuous-space version of the Wahlund effect (Wahlund 1928). Indeed, for random sampling under the spatial model, *F*_*IS*_ does not approach the random mating equivalent until neighborhood sizes of nearly 1000. On the other hand, the dependency of raw observed heterozygosity on neighborhood size is not monotone. Under midpoint sampling observed heterozygosity is inflated even over the random mating expectation, as a result of the a higher proportion of heterozygotes occurring in the middle of the landscape (Figure S6). This echoes a report from Shirk and Cushman (2014) who observed a similar excess of heterozygosity in the middle of the landscape when simulating under a lattice model.

#### IBS tracts and correlations with geographic distance

We next turn our attention to the effect of geographic distance on haplotype block length sharing, summarized for sets of nearby and distant individuals in Figure 5. There are two main patterns to note. First, nearby individuals share more long IBS tracts than distant individuals (as expected because they are on average more closely related). Second, the difference in the number of long IBS tracts between nearby and distant individuals decreases as neighborhood size increases. This reflects the faster spatial mixing of populations with higher dispersal, which breaks down the correlation between the IBS tract length distribution and geographic distance. This can also be seen in the bottom row of Figure 4B, where the correlation coefficients between the summaries of the IBS tract length distribution (the mean, skew, and count of tracts over 10^6^bp) and geographic distance approaches 0 as neighborhood size increases.

**Figure 5.**
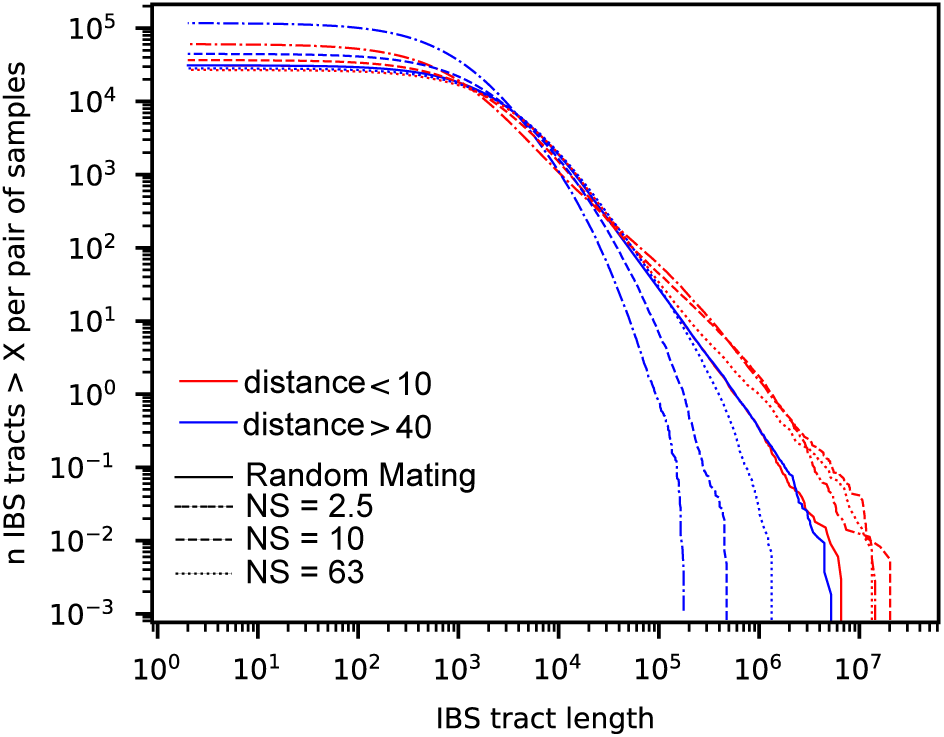
Cumulative distributions for IBS tract lengths per pair of individuals at different geographic distances, across three neighborhood sizes (NS). Nearby pairs (red curves) share many more long IBS tracts than do distant pairs (blue curves), except in the random mating model. The distribution of long IBS tracts between nearby individuals are very similar across neighborhood sizes, but distant individuals are much more likely to share long IBS tracts at high neighborhood size than at low neighborhood size.

The patterns observed for correlations of IBS tract lengths with geographic distance are similar to those observed in the more familiar correlation of allele frequency measures such as *D*_*xy*_ (i.e., “genetic distance”) or *F*_*ST*_ against geographic distance (Rousset 1997). *D*_*xy*_ is positively correlated with the geographic distance between the individuals, and the strength of this correlation declines as dispersal increases (Figure 4B), as expected (Wright 1943; Rousset 1997). This relationship is very similar across random and point sampling strategies, but is weaker for midpoint sampling, perhaps due to a dearth of long-distance comparisons. In much of empirical population genetics a regression of genetic differentiation against spatial distance is a de-facto metric of the significance of isolation by distance. The similar behavior of moments of the pairwise distribution of IBS tract lengths shows why haplotype block sharing has recently emerged as a promising source of information on spatial demography through methods described in Ringbauer *et al.* (2017) and Baharian *et al.* (2016).

#### Spatial distribution of allele copies

Mutations occur in individuals and spread geographically over time. Because low frequency alleles generally represent recent mutations (Sawyer 1977; Griffiths *et al.* 1999), the geographic spread of an allele may covary along with its frequency in the population. To visualize this relationship we calculated the average distance among individuals carrying a focal derived allele across simulations with varying neighborhood sizes, shown in Figure 6. On average we find that low frequency alleles are the most geographically restricted, and that the extent to which geography and allele frequency are related depends on the amount of dispersal in the population. For populations with large neighborhood sizes we found that even very low frequency alleles can be found across the full landscape, whereas in populations with low neighborhood sizes the relationship between distance among allele copies and their frequency is quite strong. This is the basic process underlying Novembre and Slatkin’s (2009) method for estimating dispersal distances based on the distribution of low frequency alleles, and also generates the greater degree of bias in GWAS effect sizes for low frequency alleles identified in Mathieson and McVean (2012).

**Figure 6.**
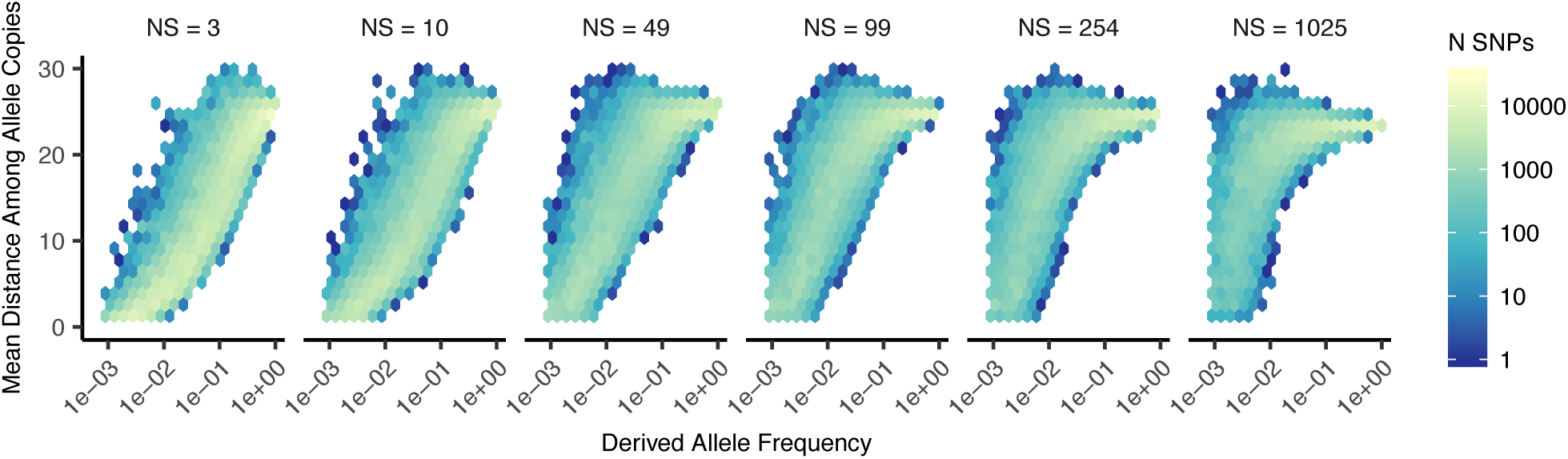
Spatial spread of rare alleles by neighborhood size (NS): Each plot shows the distribution (across derived alleles and simulations) of average pairwise distance between individuals carrying a focal derived allele and derived allele frequency.

#### Effects of Space on Demographic Inference

One of the most important uses for population genetic data is inferring demographic history of populations. As demonstrated above, the site frequency spectrum and the distribution of IBS tracts varies across neighborhood sizes and sampling strategies. Does this variation lead to different inferences of past population sizes? To ask this we inferred population size histories from samples drawn from our simulated populations with two approaches: stairwayplot (Liu and Fu 2015), which uses a genome-wide estimate of the SFS, and SMC++ (Terhorst *et al.* 2016), which incorporates information on both the SFS and linkage disequilibrium across the genome.

Figure 7A shows rolling medians of inferred population size histories from each method across all simulations, grouped by neighborhood size and sampling strategy. In general these methods tend to slightly overestimate ancient population sizes and infer recent population declines when neighborhood sizes are below 20 and sampling is spatially clustered. The overestimation of ancient population sizes however is relatively minor, averaging around a two-fold inflation at 10,000 generations before present in the worst-affected bins. For stairwayplot we found that many runs infer dramatic population bottlenecks in the last 1,000 generations when sampling is spatially concentrated, resulting in ten-fold or greater underestimates of recent population sizes. However SMC++ appeared more robust to this error, with runs on point- and midpoint-sampled simulations at the lowest neighborhood sizes underestimating recent population sizes by roughly half and those on randomly sampled simulations showing little error. Above neighborhood sizes of around 100, both methods performed relatively well when averaging across results from multiple simulations.

**Figure 7.**
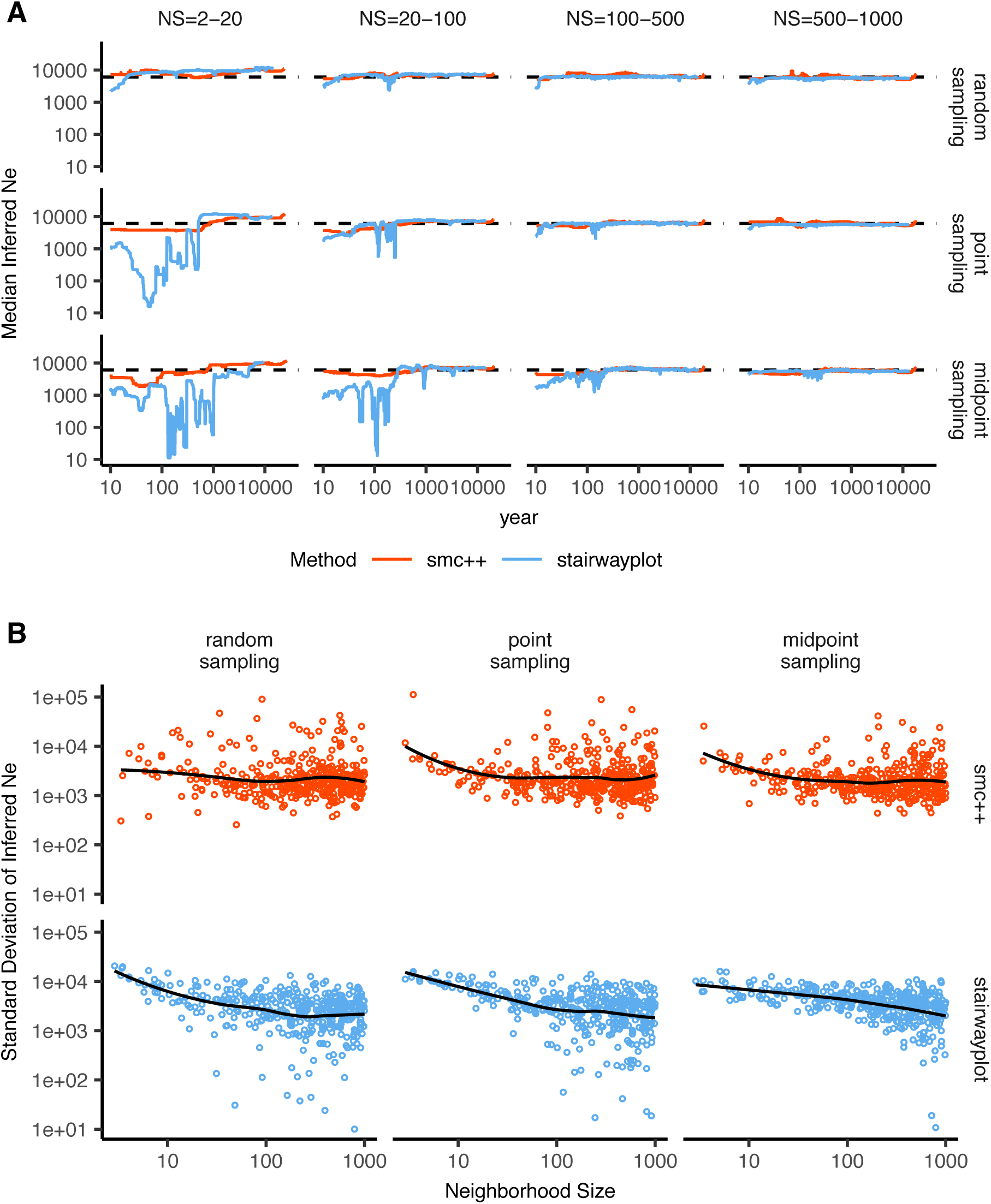
A: Rolling median inferred *N*_*e*_ trajectories for stairwayplot and smc++ across sampling strategies and neighborhood size bins. The dotted line shows the mean *N*_*e*_ of random-mating simulations. B: Standard deviation of individual inferred *N*_*e*_ trajectories, by neighborhood size and sampling strategy. Black lines are loess curves. Plots including individual model fits are shown in Figure S7.

However, individual simulations were often inferred to have turbulent demographic histories, as shown by the individually inferred histories (shown in Figure S7). Indeed, the standard deviation of inferred *N*_*e*_ across time points (shown in Figure 7B) often exceeds the expected *N*_*e*_ for both methods. That is, despite the nearly constant population sizes in our simulations, both methods tended to infer large fluctuations in population size over time, which could potentially result in incorrect biological interpretations. On average the variance of inferred population sizes was elevated at the lowest neighborhood sizes and declines as dispersal increases, with the strongest effects seen in stairwayplot results with clustered sampling and neighborhood sizes less than 20 (Figure 7B).

### GWAS

To ask what confounding effects spatial genetic variation might have on genome-wide association studies we performed GWAS on our simulations using phenotypes that were determined solely by the environment – so, any SNP showing statistically significant correlation with phenotype is a false positive. As expected, spatial autocorrelation in the environment causes spurious associations across much of the genome if no correction for genetic relatedness among samples is performed (Figures 8 and S8). This effect is particularly strong for clinal and corner environments, for which the lowest dispersal levels cause over 60% of SNPs in the sample to return significant associations. Patchy environmental distributions, which are less strongly spatially correlated (Figure 8A), cause fewer false positives overall but still produce spurious associations at roughly 10% of sites at the lowest neighborhood sizes. Interestingly we also observed a small number of false positives in roughly 3% of analyses on simulations with nonspatial environments, both with and without PC covariates included in the regression.

**Figure 8.**
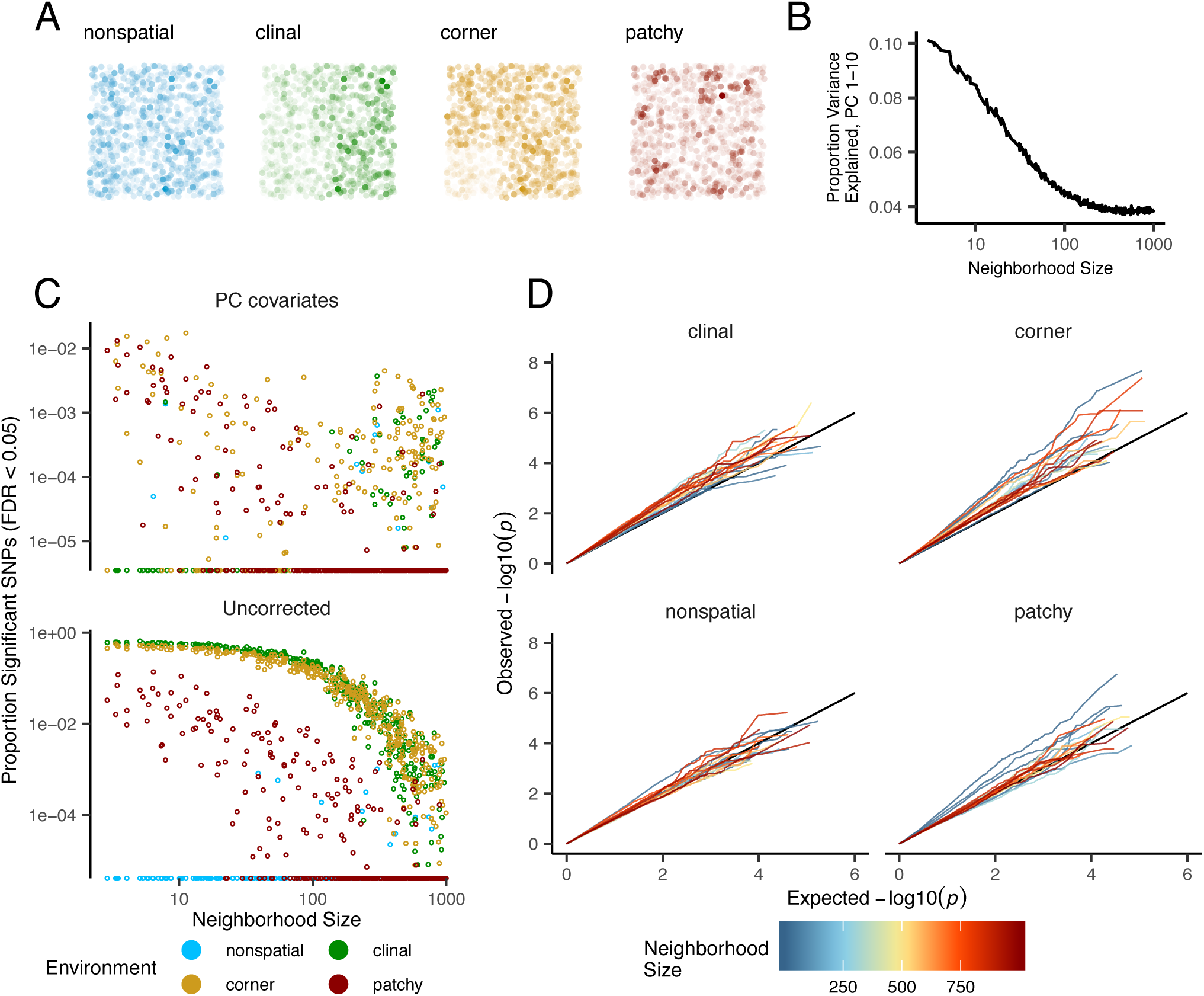
Impacts of spatially varying environments and isolation by distance on linear regression GWAS. Simulated quantitative phenotypes are determined only by an individual’s location and the spatial distribution of environmental factors. In **A** we show the phenotypes and locations of sampled individuals under four environmental distributions, with transparency scaled to phenotype. As neighborhood size increases a PCA explains less of the total variation in the data (**B**). Spatially correlated environmental factors cause false positives at a large proportion of SNPs, which is partially but not entirely corrected by adding the first 10 PC coordinates as covariates (**C**). Quantile-quantile plots in (**D**) show inflation of − log_10_(*p*) after PC correction for simulations with spatially structured environments, with line colors showing the neighborhood size of each simulation.

The confounding effects of geographic structure are well known, and it is common practice to control for this by including principal components (PCs) as covariates to control for these effects. This mostly works in our simulations – after incorporating the first ten PC axes as covariates, the vast majority of SNPs no longer surpass a significance threshold chosen to have a 5% false discovery rate (FDR). However, a substantial number of SNPs – up to 1.5% at the lowest dispersal distances – still surpass this threshold (and thus would be false positives in a GWAS), especially under “corner” and “patchy” environmental distributions (Figure 8C). At neighborhood sizes larger than 500, up to 0.31% of SNPs were significant for corner and clinal environments. Given an average of 132,000 SNPs across simulations after MAF filtering, this translates to up to 382 false-positive associations; for human-sized genomes, this number would be much larger. In most cases the *p* values for these associations were significant after FDR correction but would not pass the threshold for significance under the more conservative Bonferroni correction (see example Manhattan plots in figure S8).

Clinal environments cause an interesting pattern in false positives after PC correction: at low neighborhood sizes the correction removes nearly all significant associations, but at neighborhood sizes above roughly 250 the proportion of significant SNPs increases to up to 0.4% (Figure 8). This may be due to a loss of descriptive power of the PCs – as neighborhood size increases, the total proportion of variance explained by the first 10 PC axes declines from roughly 10% to 4% (Figure 8B). Essentially, PCA seems unable to effectively summarize the weak population structure present in large-neighborhood simulations, but these populations continue to have enough spatial structure to create significant correlations between genotypes and the environment. A similar process can also be seen in the corner phenotype distribution, in which the count of significant SNPs initially declines as neighborhood size increases and then increases at approximately the point at which the proportion of variance explained by PCA approaches its minimum.

Figure 8D shows quantile-quantile plots for a subset of simulations that show the degree of genome-wide inflation of test statistics in PC-corrected GWAS across all simulations and environmental distributions. An alternate visualization is also included in figure S9. For clinal environments, − log_10_(*p*) values are most inflated when neighborhood sizes are large, consistent with the pattern observed in the count of significant associations after PC regression. In contrast corner and patchy environments cause the greatest inflation in −log_10_(*p*) at neighborhood sizes less than 100, which likely reflects the inability of PCA to account for fine-scale structure caused by very limited dispersal. Finally, we observed that PC regression appears to overfit to some degree for all phenotype distributions, visible in Figure 8D as points falling below the 1:1 line.

## Discussion

In this study, we have used efficient forward time population genetic simulations to describe the myriad influence of continuous geography on genetic variation. In particular, we examine how three main types of downstream empirical inference are affected by unmodeled spatial population structure – population genetic summary statistics, inference of population size history, and genome-wide association studies (GWAS). As discussed above, space often matters (and sometimes dramatically), both because of how samples are arranged in space, and because of the inherent patterns of relatedness established by geography.

### Effects of Dispersal

Limited dispersal inflates effective population size, creates correlations between genetic and spatial distances, and introduces strong distortions in the site frequency spectrum that are reflected in a positive Tajima’s *D* (Figure 4). At the lowest dispersal distances, this can increase genetic diversity threefold relative to random-mating expectations. These effects are strongest when neighborhood sizes are below 100, but in combination with the effects of nonrandom sampling they can persist up to neighborhood sizes of at least 1000 (e.g., inflation in Tajima’s *D* and observed heterozygosity under midpoint sampling). If samples are chosen uniformly from across space, the general pattern is similar to expectations of the original analytic model of Wright (1943), which predicts that populations with neighborhood sizes under 100 will differ substantially from random mating, while those above 10,000 will be nearly indistinguishable from panmixia.

The patterns observed in sequence data reflect the effects of space on the underlying genealogy. Nearby individuals coalesce rapidly under limited dispersal and so are connected by short branch lengths, while distant individuals take much longer to coalesce than they would under random mating. Mutation and recombination events in our simulation both occur at a constant rate along branches of the genealogy, so the genetic distance and number of recombination events separating sampled individuals simply gives a noisy picture of the genealogies connecting them. Tip branches (i.e., branches subtending only one individual) are then relatively short, and branches in the middle of the genealogy connecting local groups of individuals relatively long, leading to the biases in the site frequency spectrum shown in Figure 4.

The genealogical patterns introduced by limited dispersal are particularly apparent in the distribution of haplotype block lengths (Figure 4). This is because identical-by-state tract lengths reflect the impacts of two processes acting along the branches of the underlying genealogy – both mutation and recombination – rather than just mutation as is the case when looking at the site frequency spectrum or related summaries. This means that the pairwise distribution of haplotype block lengths carries with it important information about genealogical variation in the population, and correlation coefficients between moments of the this distribution and geographic location contain signal similar to the correlations between *F*_*ST*_ or *D*_*xy*_ and geographic distance (Rousset 1997). Indeed this basic logic underlies two recent studies explicitly estimating dispersal from the distribution of shared haplotype block lengths (Ringbauer *et al.* 2017; Baharian *et al.* 2016). Conversely, because haplotype-based measures of demography are particularly sensitive to variation in the underlying genealogy, inference approaches that assume random mating when analyzing the distribution of shared haplotype block lengths are likely to be strongly affected by spatial processes.

### Effects of Sampling

One of the most important differences between random mating and spatial models is the effect of sampling: in a randomly mating population the spatial distribution of sampling effort has no effect on estimates of genetic variation (Table S1), but when dispersal is limited sampling strategy can compound spatial patterns in the underlying genealogy and create pervasive impacts on all downstream genetic analyses (see also Städler *et al.* (2009)). In most species, the difficulty of traveling through all parts of a species range and the inefficiency of collecting single individuals at each sampling site means that most studies follow something closest to the “point” sampling strategy we simulated, in which multiple individuals are sampled from nearby points on the landscape. For example, in ornithology a sample of 10 individuals per species per locality is a common target when collecting for natural history museums. In classical studies of *Drosophila* variation the situation is considerably worse, in which a single orchard might be extensively sampled.

When sampling is clustered at points on a landscape and dispersal is limited, the sampled individuals will be more closely related than a random set of individuals. Average coalescence times of individuals collected at a locality will then be more recent and branch lengths shorter than expected by analyses assuming random mating. This leads to fewer mutations and recombination events occurring since their last common ancestor, causing a random set of individuals to share longer average IBS tracts and have fewer nucleotide differences. For some data summaries, such as Tajima’s *D*, Watterson’s *θ*, or the correlation coefficient between spatial distance and the count of long haplotype blocks, this can result in large differences in estimates between random and point sampling (Figure 4). Inferring underlying demographic parameters from these summary statistics – unless the spatial locations of the sampled individuals are somehow taken into account – will likely be subject to bias.

We observed the largest sampling effects using “midpoint” sampling. This model is meant to reflect a bias in sampling effort towards the middle of a species’ range. In empirical studies this sampling strategy could arise if, for example, researchers choose to sample the center of the range and avoid range edges to maximize probability of locating individuals during a short field season. Because midpoint sampling provides limited spatial resolution it dramatically reduces the magnitude of observed correlations between spatial and genetic distances. More surprisingly, midpoint sampling also leads to strongly positive Tajima’s *D* and an inflation in the proportion of heterozygous individuals in the sample – similar to the effect of sampling a single deme in an island model as reported in Städler *et al.* (2009). This increase in observed heterozygosity appears to reflect the effects of range edges, which are a fundamental facet of spatial genetic variation. If individuals move randomly in a finite two-dimensional landscape then regions in the middle of the landscape receive migrants from all directions while those on the edge receive no migrants from at least one direction. The average number of new mutations moving into the middle of the landscape is then higher than the number moving into regions near the range edge, leading to higher heterozygosity and lower inbreeding coefficients (*F*_*IS*_) away from range edges. Though here we used only a single parameterization of fitness decline at range edges we believe this is a general property of non-infinite landscapes as it has also been observed in previous studies simulating under lattice models (Neel *et al.* 2013; Shirk and Cushman 2014).

In summary, we recommend that empirical researchers collect individuals from across as much of the species’ range as practical, choosing samples separated by a range of spatial scales. Many summary statistics are designed for well-mixed populations, and so provide different insights into genetic variation when applied to different subsets of the population. Applied to a cluster of samples, summary statistics based on segregating sites (e.g., Watterson’s *θ* and Tajima’s *D*), heterozygosity, or the distribution of long haplotype blocks, can be expected to depart significantly from what would be obtained from a wider distribution of samples. Comparing the results of analyses conducted on all individuals versus those limited to single individuals per locality can provide an informative contrast. Finally we wish to point out that the bias towards intermediate allele frequencies that we observe may mean that the importance of linked selection, at least as is gleaned from the site frequency spectrum, may be systematically underestimated currently.

### Demography

Previous studies have found that population structure and nonrandom sampling can create spurious signals of population bottlenecks when attempting to infer demographic history with microsatellite variation, summary statistics, or runs of homozygosity (Chikhi *et al.* 2010; Städler *et al.* 2009; Ptak and Przeworski 2002; Mazet *et al.* 2015). Here we found that methods that infer detailed population trajectories through time based on the SFS and patterns of LD across the genome are also subject to this bias, with some combinations of dispersal and sampling strategy systematically inferring deep recent population bottlenecks and overestimating ancient *N*_*e*_ by around a factor of 2. We were surprised to see that both stairwayplot and SMC++ can tolerate relatively strong isolation by distance – i.e., neighborhood sizes of 20 – and still perform well when averaging results across multiple simulations. Inference in populations with neighborhood sizes over 20 was relatively unbiased unless samples were concentrated in the middle of the range (Figure 7). Although median demography estimates across many independent simulations were fairly accurate, empirical work has only a single estimate to work with, and individual model fits (Figure S7) suggest that spuriously inferred population size changes and bottlenecks are common, especially at small neighborhood sizes. As we will discuss below, most empirical estimates of neighborhood size, including all estimates for human populations, are large enough that population size trajectories inferred by these approaches should not be strongly affected by spatial biases created by dispersal in continuous landscapes. In contrast, Mazet *et al.* (2015) found that varying migration rates through time could create strong biases in inferred population trajectories from an *n*-island model with parameters relevant for human history, suggesting that changes in migration rates through time are more likely to drive variation in inferred *N*_*e*_ than isolation by distance.

We found that SMC++ was more robust to the effects of space than stairwayplot, underestimating recent populations by roughly half in the worst time periods rather than nearly 10-fold as with stairwayplot. Though this degree of variation in population size is certainly meaningful in an ecological context, it is relatively minor in population genetic terms. Methods directly assessing haplotype structure in phased data example, (e.g., Browning and Browning 2015) are thought to provide increased resolution for recent demographic events, but in this case the error we observed was essentially an accurate reflection of underlying genealogies in which terminal branches are anomalously short. Combined with our analysis of IBS tract length variation (Figure 5) this suggests that haplotype-based methods are likely to be affected by similar biases.

A more worrying pattern was the high level of variance in inferred *N*_*e*_ trajectories for individual model fits using these methods, which was highest in simulations with the smallest neighborhood size (Figure 7, Figure S7). This suggests that, at a minimum, researchers working with empirical data should replicate analyses multiple times and take a rolling average if model fits are inconsistent across runs. Splitting samples and running replicates on separate subsets – the closest an empirical study can come to our design of averaging the results from multiple simulations – may also alleviate this issue.

Our analysis suggests that many empirical analyses of population size history using methods like SMC++ are robust to error caused by spatial structure within continuous landscapes. Inferences drawn from static SFS-based methods like stairwayplot should be treated with caution when there are signs of isolation by distance in the underlying data (for example, if a regression of *F*_*ST*_ against the logarithm of geographic distance has a significantly positive slope), and in particular an inference of population bottlenecks in the last 1000 years should be discounted if sampling is clustered, but estimates of deeper time patterns are likely to be fairly accurate. The biases in the SFS and haplotype structure identified above (see also Wakeley 1999; Chikhi *et al.* 2010; Städler *et al.* 2009) are apparently small enough that they fall within the range of variability regularly inferred by these approaches, at least on datasets of the size we simulated.

### GWAS

Spatial structure is particularly challenging for genome-wide association studies, because the effects of dispersal on genetic variation are compounded by spatial variation in the environment (Mathieson and McVean 2012). Spatially restricted mate choice and dispersal causes variation in allele frequencies across the range of a species. If environmental factors affecting the phenotype of interest also vary over space, then allele frequencies and environmental exposures will covary over space. In this scenario an uncorrected GWAS will infer genetic associations with a purely environmental phenotype at any site in the genome that is differentiated over space, and the relative degree of bias will be a function of the degree of covariation in allele frequencies and the environment (i.e., Figure 8C, bottom panel). This pattern has been demonstrated in a variety of simulation and empirical contexts (Price *et al.* 2006; Yu *et al.* 2005; Young *et al.* 2018; Mathieson and McVean 2012; Kang *et al.* 2008, 2010; Bulik-Sullivan *et al.* 2015; Berg *et al.* 2018; Sohail *et al.* 2018).

Incorporating PC positions as covariates in a linear-regression GWAS (Price *et al.* 2006) is designed to address this challenge by regressing out a baseline level of “average” differentiation. In essence, a PC-corrected GWAS asks “what regions of the genome are more associated with this phenotype than the average genome-wide association observed across populations?” In our simulations, we observed that this procedure can fail under a variety of circumstances. If dispersal is limited and environmental variation is clustered in space (i.e., corner or patchy distributions in our simulations), PC positions fail to capture the fine-scale spatial structure required to remove all signals of association. Conversely, as dispersal increases, PCA loses power to describe population structure before spatial mixing breaks down the relationship between genotype and the environment. These effects were observed with all spatially correlated environmental patterns, but were particularly pronounced if environmental effects are concentrated in one region, as was also found by Mathieson and McVean (2012). Though increasing the number of PC axes used in the analysis may reduce the false-positive rate, this may also decrease the power of the test to detect truly causal alleles (Lawson *et al.* 2019).

In this work we simulated a single chromosome with size roughly comparable to one human chromosome. If we scale the number of false-positive associations identified in our analyses to a GWAS conducted on whole-genome data from humans, we would expect to see several thousand weak false-positive associations after PC corrections in a population with neighborhood sizes up to at least 1000 (which should include values appropriate for many human populations). Notably, very few of the spurious associations we identified would be significant at a conservative Bonferroni-adjusted *p*-value cutoff (see Figure S8). This suggests that GWAS focused on finding strongly associated alleles for traits controlled by a limited number of variants in the genome are likely robust to the impacts of continuous spatial structure. However, methods that analyze the combined effects of thousands or millions of weakly associated variants such as polygenic risk scores (Khera *et al.* 2018) are likely to be affected by subtle population structure. Indeed as recently identified in studies of genotype associations for human height in Europe (Berg *et al.* 2018; Sohail *et al.* 2018), PC regression GWAS in modern human populations do include residual signal of population structure in large-scale analyses of polygenic traits. When attempting to make predictions across populations with different environmental exposures, polygenic risk scores affected by population structure can be expected to offer low predictive power, as was shown in a recent study finding lower performance outside European populations (Martin *et al.* 2019).

In summary, spatial covariation in population structure and the environment confounds the interpretation of GWAS *p*-values, and correction using principal components is insufficient to fully separate these signals for polygenic traits under a variety of environmental and population parameter regimes. Other GWAS methods such as mixed models (Kang *et al.* 2008) may be less sensitive to this confounding, but there is no obvious reason that this should be so. One approach to estimating the degree of bias in GWAS caused by population structure is LD score regression (Bulik-Sullivan *et al.* 2015). Though this approach appears to work well in practice, its interpretation is not always straightforward and it is likely biased by the presence of linked selection (Berg *et al.* 2018). In addition, we observed that in many cases the false-positive SNPs we identified appeared to be concentrated in LD peaks similar to those expected from truly causal sites (Figure S8), which may confound LD score regression.

We suggest a straightforward alternative for species in which the primary axes of population differentiation is space (note this is likely not the case for some modern human populations): run a GWAS with spatial coordinates as phenotypes and check for *p*-value inflation or significant associations. If significant associations with sample locality are observed after correcting for population structure, the method is sensitive to false positives induced by spatial structure. This is essentially the approach taken in our “clinal” model (though we add normally distributed noise to our phenotypes). This approach has recently been taken with polygenic scores for UK Biobank samples in Haworth *et al.* (2019), finding that scores are correlated with birth location even in this relatively homogenous sample. Of course, it is possible that genotypes indirectly affect individual locations by adjusting organismal fitness and thus habitat selection across spatially varying environments, but we believe that this hypothesis should be tested against a null of stratification bias inflation rather than accepted as true based on GWAS results.

### Where are natural populations on this spectrum?

For how much of the tree of life do spatial patterns circumscribe genomic variation? In Table 1 we gathered estimates of neighborhood size from a range of organisms to get an idea of how strongly local geographic dispersal affects patterns of variation. This is an imperfect measure: some aspects of genetic variation are most strongly determined by neighborhood size (Wright 1946), others (e.g., number of segregating sites) are more strongly determined by global *N*_*e*_ or by the ratio of the two. In addition, these empirical examples are likely biased towards small-neighborhood species (because few studies have quantified neighborhood size in species with very high dispersal or population density). However, from the available data we find that neighborhood sizes in the range we simulated are fairly common across a range of taxa. At the extreme low end of empirical neighborhood size estimates we see some flowering plants, large mammals, and colonial insects like ants. Species such as this have neighborhood size estimates small enough that spatial processes are likely to strongly influence inference. These include some human populations such as the Gainj- and Kalam-speaking people of Papua New Guinea, in which the estimated neighborhood sizes in Rousset (1997) range from 40 to 410 depending on the method of estimation. Many more species occur in a middle range of neighborhood sizes between 100 and 1000 – a range in which spatial processes play a minor role in our analyses under random spatial sampling but are important when sampling of individuals in space is clustered. Surprisingly, even some flying insects with huge census population sizes fall in this group, including fruit flies (*D. melanogaster*) and mosquitoes (*A. aegypti*). Last, many species likely have neighborhood sizes much larger than we simulated, including the recent ancestors of modern humans in northeastern Europe (Ringbauer *et al.* 2017). For these species demographic inference and summary statistics are likely to reflect minimal bias from spatial effects as long as dispersal is truly continuous across the landscape. While that is so we caution that association studies in which the effects of population structure are confounded with spatial variation in the environment are still sensitive to dispersal even at these large neighborhood sizes.

### Other demographic models

Any simulation of a population of reproducing organisms requires some kind of control on population sizes, or else the population will either die out or grow very large after a sufficiently long period of time. The usual choice of population regulation for population genetics – a constant size, as in the Wright–Fisher model – implies biologically unrealistic interactions between geographically distant parts of the species range. Our choice to regulate population size by including a local density-dependent control on mortality is only one of many possible ways to do this. We could have instead regulated fecundity, or recruitment, or both; this general class of models is sometimes referred to as the “Bolker–Pacala model” (Bolker and Pacala 1997). It is not currently clear how much different choices of demographic parameters, or of functional forms for the regulation, might quantitatively affect our results, although the general predictions should be robust to similar forms of regulation. Since populations are still entirely *intrinsically* regulated, our model still has a very strong “population genetics” flavor. Alternatively, population size could be regulated by interactions with other species (e.g., a Lotka-Volterra model), or extrinsically specified by local resource availability (e.g., by food or nest site availability). Indeed, our model could be interpreted as a caricature of such a model: as local density increases, good habitat is increasingly occupied, pushing individuals into more marginal habitat and increasing their mortality. Many such models should behave similarly to ours, but others (especially those with local population cycling), may differ dramatically.

Population genetic simulations often use grids of discrete demes, which are assumed to approximate continuous space. However, there are theoretical reasons to expect that increasingly fine grids of discrete demes do not approach the continuous model (Barton *et al.* 2002). If continuous space can be approximated by a limit of discrete models, this should be true regardless of the precise details of the discrete model. Although we carefully chose parameters to match our continuous models, we found that some aspects of genetic variation diverged from the continuous case as the discretization got finer. This suggests that these models do not converge in the limit. However, many populations may indeed be well-modeled as a series of discrete, randomly-mating demes if, for example, suitable habitats are patchily distributed across the landscape. There is a clear need for greater exploration of the consequences for population genetics of ecologically realistic population models.

### Future Directions and Limitations

As we have shown, a large number of population genetic summary statistics contain information about spatial population processes. We imagine that combinations of such summaries might be sufficient for the construction of supervised machine learning regressors (e.g., Schrider and Kern 2018) for the accurate estimation of dispersal from genetic data. Indeed, Ashander *et al.* (2018) found that inverse interpolation on a vector of summary statistics provided a powerful method of estimating dispersal distances. Expanding this approach to include the haplotype-based summary statistics studied here and applying machine learning regressors built for general inference of nonlinear relationships from high-dimensional data may allow precise estimation of spatial parameters under a range of complex models.

One facet of spatial variation that we did not address in this study is the confounding of dispersal and population density implicit in the definition of Wright’s neighborhood size. Our simulations were run under constant densities, but Guindon *et al.* (2016) and Ringbauer *et al.* (2017) have shown that these parameters are identifiable under some continuous models. Similarly, though the scaling effects of dispersal we show in Figure 4 should occur in populations of any total size, other aspects such as the number of segregating sites are also likely affected by the total landscape size (and so total census size). Much additional work remains to be done to better understand how these parameters interact to shape genetic variation in continuous space, which we leave to future studies.

Though our simulation allows incorporation of realistic demographic and spatial processes, it is inevitably limited by the computational burden of tracking tens or hundreds of thousands of individuals in every generation. In particular, computations required for mate selection and spatial competition scale approximately with the product of the total census size and the neighborhood size and so increase rapidly for large populations and dispersal distances. The reverse-time spatial Lambda–Fleming–Viot model described by Barton *et al.* (2010) and implemented by Kelleher *et al.* (2014) allows exploration of larger population and landscape sizes, but the precise connection of these models to forward-time demography is not yet clear. Alternatively, implementation of parallelized calculations may allow progress with forward-time simulations.

Finally, we believe that the difficulties in correcting for population structure in continuous populations using principal components analysis or similar decompositions is a difficult issue, well worth considering on its own. How can we best avoid spurious correlations while correlating genetic and phenotypic variation without underpowering the methods? Perhaps optimistically, we posit that process-driven descriptions of ancestry and/or more generalized unsupervised methods may be able to better account for carry out this task.

## Data Availability

Scripts used for all analyses and figures are available at https://github.com/kern-lab/spaceness.

## Acknowledgements

We thank Brandon Cooper, Matt Hahn, Doc Edge, and others for reading and thinking about this manuscript. CJB and ADK were supported by NIH award R01GM117241.

## Comparisons with Stepping-Stone Models

We also compared our model results to a regular grid of discrete populations, which is commonly used as an approximation of continuous geography. An important reason that this approximation is often made is that it allows more efficient, coalescent simulations; we implemented these using `msprime` (Kelleher *et al.* 2016). In this class of models we imagine an *n* × *n* grid of populations exchanging migrants with neighboring populations at rate *m*. If these models are good approximations of the continuous case we expect that results will converge as *n* → ∞ (while scaling *m* appropriately and keeping total population size fixed), so we ran simulations while varying *n* from 5 to 50 (Table A1). To compare with continuous models we first distributed the the same “effective” number of individuals across the landscape as in our continuous-space simulations (≈ 6100, estimated from *θ*_*π*_ of random-mating continuous-space simulations). We then approximate the mean per-generation dispersal distance *σ* given a total landscape width *W* as the product of the probability of an individual being a migrant and the distance traveled by migrants: *σ* = 4*m*(*W*/*n*). This means that *m* in different simulations with the same *σ* scales with 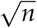. We ran 500 simulations for each value of *n* while sampling *σ* from *U*(0.2, 4). We then randomly selected 60 diploid individuals from each simulation (approximating diploidy by combining pairs of chromosomes with contiguous indices within demes) and calculated a set of six summary statistics using the scripts described in the summary statistics portion of the main text.

**Table A1.**
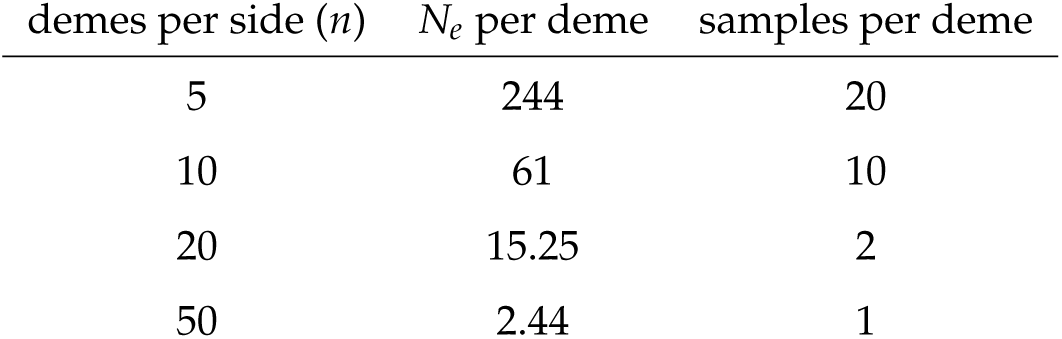
stepping-stone simulation parameters.

| demes per side ($n$) | $N_e$ per deme | samples per deme |
| --- | --- | --- |
| 5 | 244 | 20 |
| 10 | 61 | 10 |
| 20 | 15.25 | 2 |
| 50 | 2.44 | 1 |

**Figure A1.**
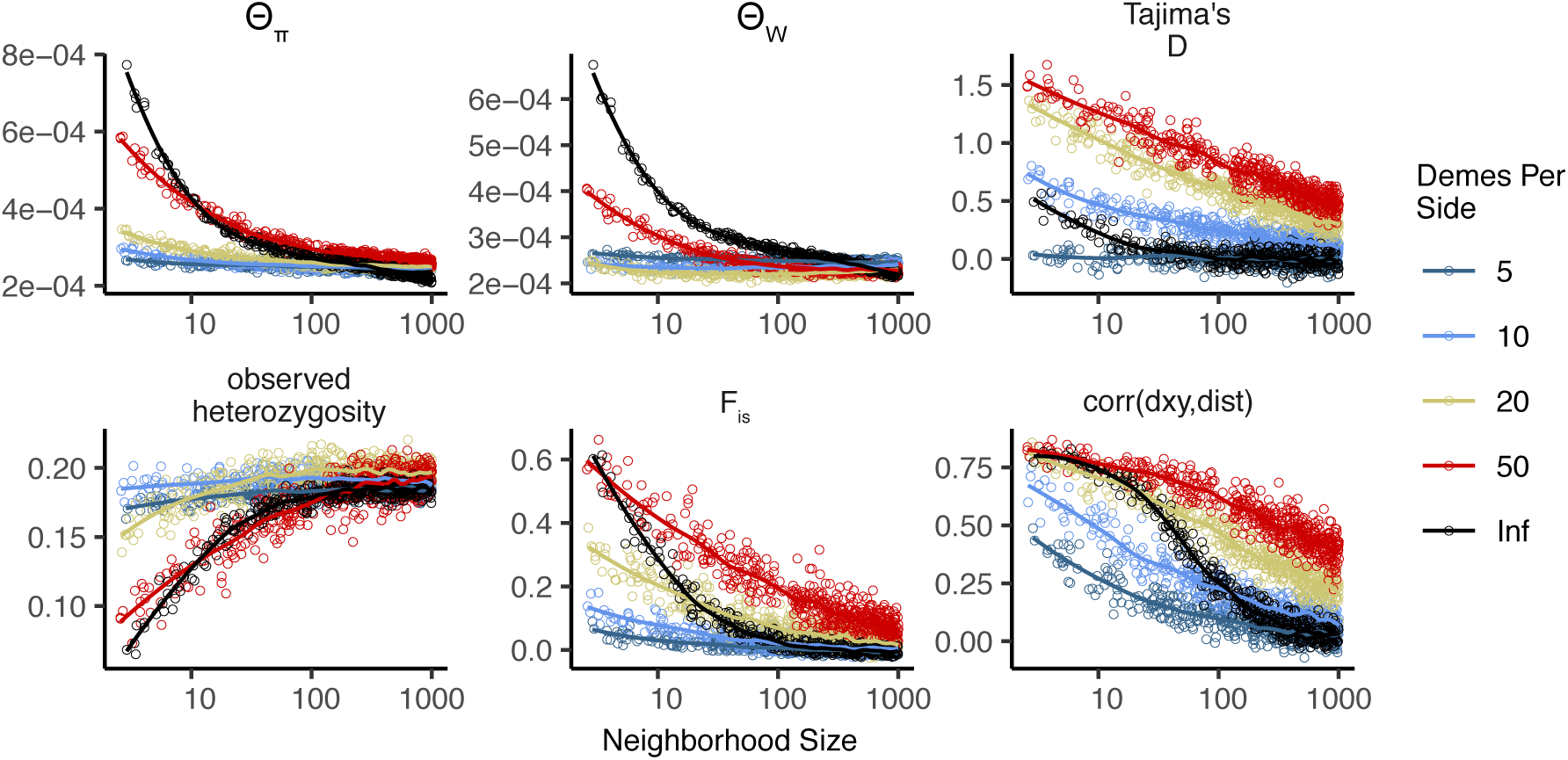
Summary statistics for 2-dimensional coalescent stepping-stone models with fixed total *N*_*e*_ and varying numbers of demes per side. The black “infinite” points are from our forward-time continuous space model. Inter-deme migration rates are related to *σ* as described above.

In general we find many of the qualitative trends are similar among continuous and stepping-stone models and that, at low neighborhood sizes, many (but not all) statistics from stepping-stone models approach the continuous model as the resolution of the grid increases. For example, *θ*_*π*_ is inflated at low neighborhood sizes (i.e., low *m*), and the extent of the inflation increases to approach the continuous case as the resolution of the landscape increases. Similar patterns are observed for *F*_*IS*_ and observed heterozygosity. However, *θ*_*W*_ behaves differently, showing a non-monotonic relationship with grid resolution. This results in an increasingly positive Tajima’s *D* in grid simulations at small neighborhood sizes, to a much greater extent than seen in a continuous model. In contrast to *θ*_*π*_, increasing the resolution of the grid causes Tajima’s *D* to deviate *more* from what is seen in the continuous model.

These differences relative to our continuous model mainly reflect two shortcomings of the reverse-time stepping stone model. If we simulate a coarse grid with relatively large populations in each deme, we cannot accurately capture the dynamics of small neighborhood sizes because mating within each deme remains random regardless of the migration rate connecting demes. This likely explains the trends in *θ*_*π*_, observed heterozygosity, and *F*_*IS*_. However increasing the number of demes while holding the total number of individuals constant results in small within-deme populations for which even the minimum sample size of 1 approaches the local *N*_*e*_ (Table A1). This results in an excess of short terminal branches in the coalescent tree, which decreases the total branch length and leads to fewer segregating sites, deflated *θ*_*W*_, and inflated Tajima’s *D*. Overall, the stepping-stone model reproduces important features of spatial structure in our continuous space model, such as a decline in *θ*_*π*_ and correlations between spatial and genetic distance with increasing migration, but introduces artifacts caused by binning the landscape into discrete demes.

## Demographic model

Local population regulation is controlled by two parameters, *L*, and *K*. Here, we show that these should be close to the average lifespan of an individual and the average number of individuals per unit area, respectively. We chose our demographic model so that every individual has on average 1/*L* offspring each time step, and if the local population density of an individual is *n*, then their probability of survival until the next time step is (equation (1)):

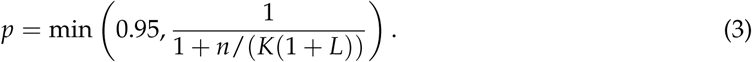

We capped survival at 0.95 so that we would not have exceptionally long-lived individuals in sparsely populated areas – otherwise, an isolated individual might live for a very long time. Since 1 *p* − ≈ *n*/(*K*(1 + *L*)), mortality goes up roughly linearly with the number of neighbors (on a scale given by *K*), as would be obtained if, for instance, mortality is due to agonistic interactions. Ignoring migration, a region is at demographic equilibrium if the per-capita probability of death is equal to the birth rate, i.e., if 1 − *p* = 1/*L*. (Note that there is no effect of age in the model, which would make the analysis more complicated.) Solving this for *n*, we get that in a well-mixed population, the equilibrium density should be around

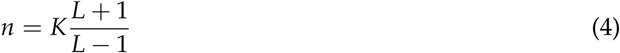

individuals per unit area. At this density, the per-capita death rate is 1/*L*, so the mean lifetime is *L*. This equilibrium density is *not K*, but (since *L* = 4) is two-thirds larger. However, in practice this model leads to a total population size which is around *K* multiplied by total geographic area (but which depends on *σ*, as discussed above). The main reason for this is that since offspring tend to be near their parents, individuals tend to be “clumped”, and so experience a higher average density than the “density” one would compute by dividing census size by geographic area (Lloyd 1967). To maintain a constant expected total population size would require making (say) *K* depend on *σ*; however, typical local population densities might then be more dissimilar.

## Supplementary Figures and Tables

**Figure S1.**
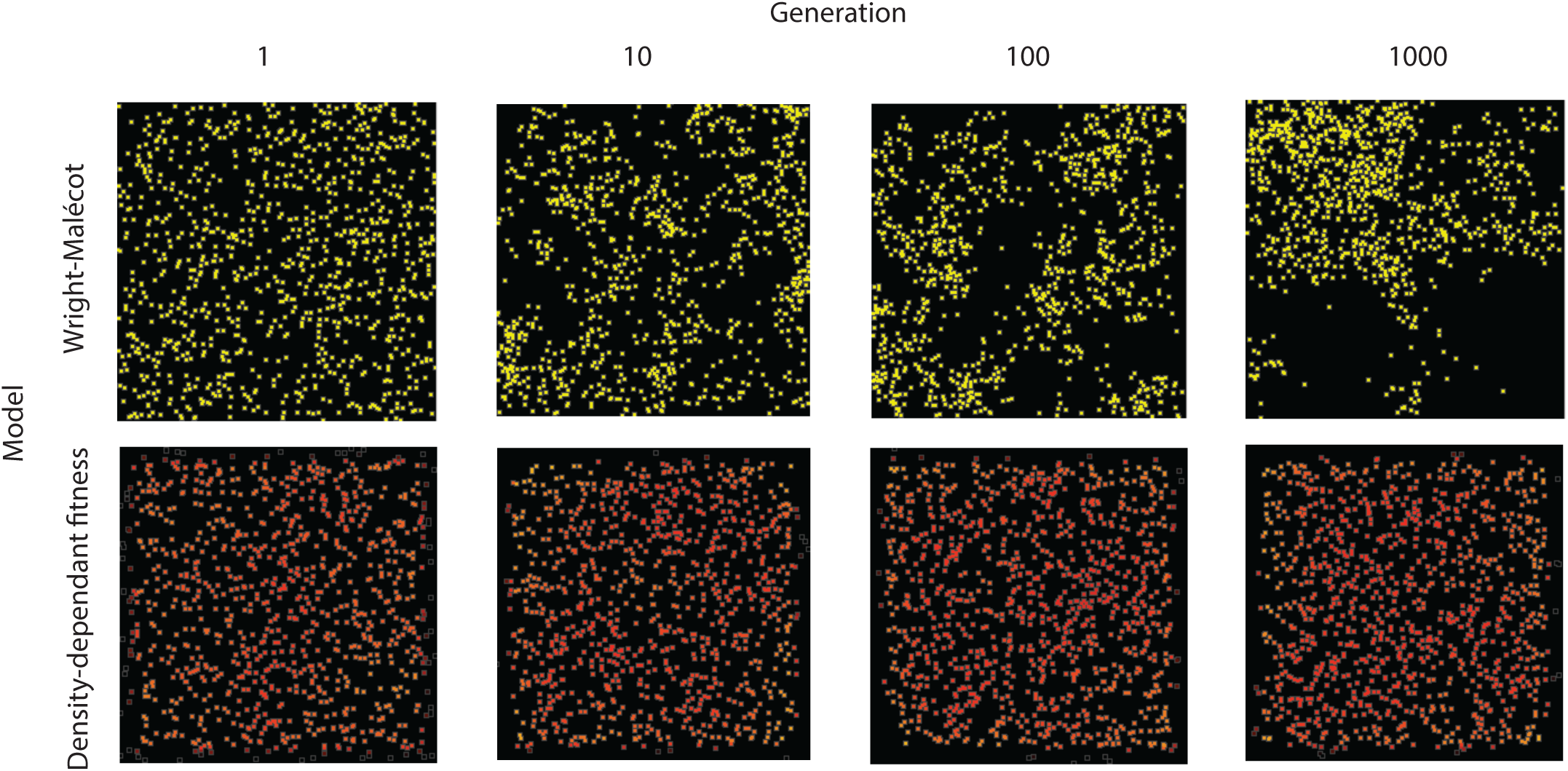
Maps of individual locations in a continuous-space Wright-Malécot model with independent dispersal of all individuals (top) and under our continuous space model incorporating density-dependant fitness (bottom). The clustering seen in the top row is the “Pain in the Torus” described by Felsenstein (1975).

**Figure S2.**
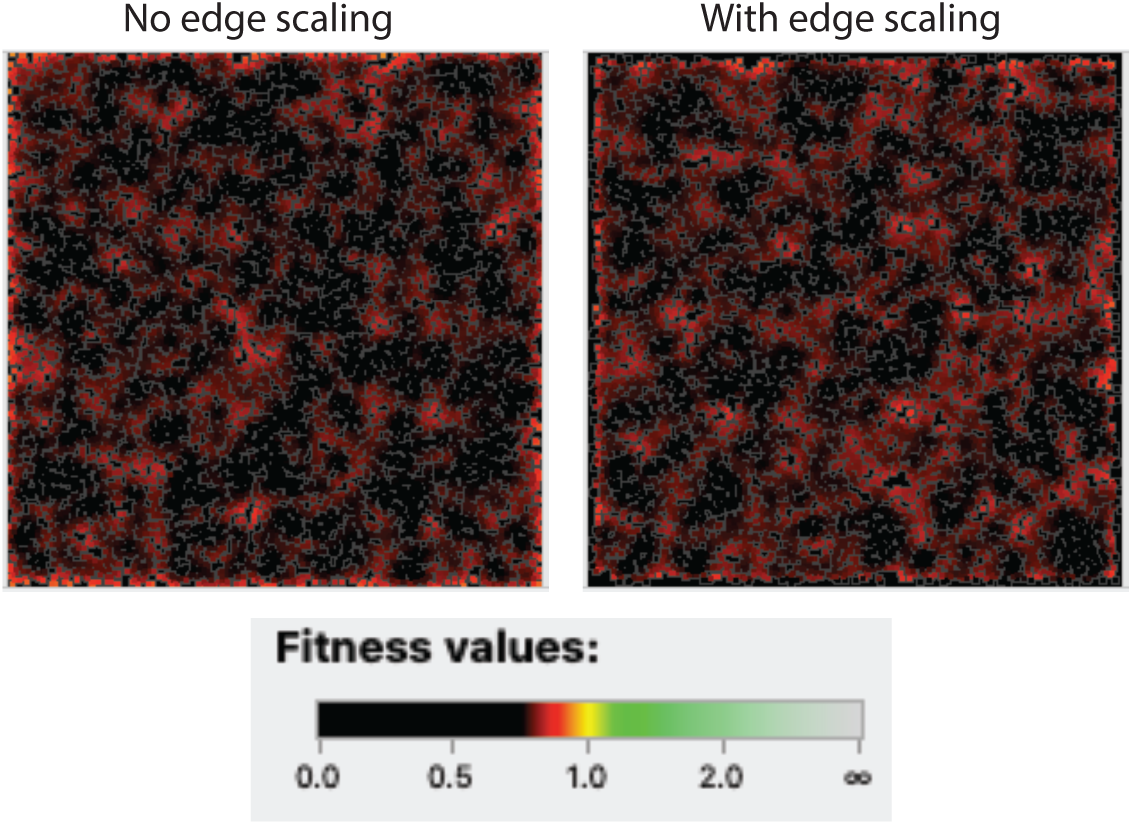
Comparison of individual fitness across the landscape in simulations with (right) and without (left) a decline in fitness approaching range edges. Note the slight excess of high-fitness individuals at edges on the left, which is (partially) counteracted by the scaling procedure.

**Figure S3.**
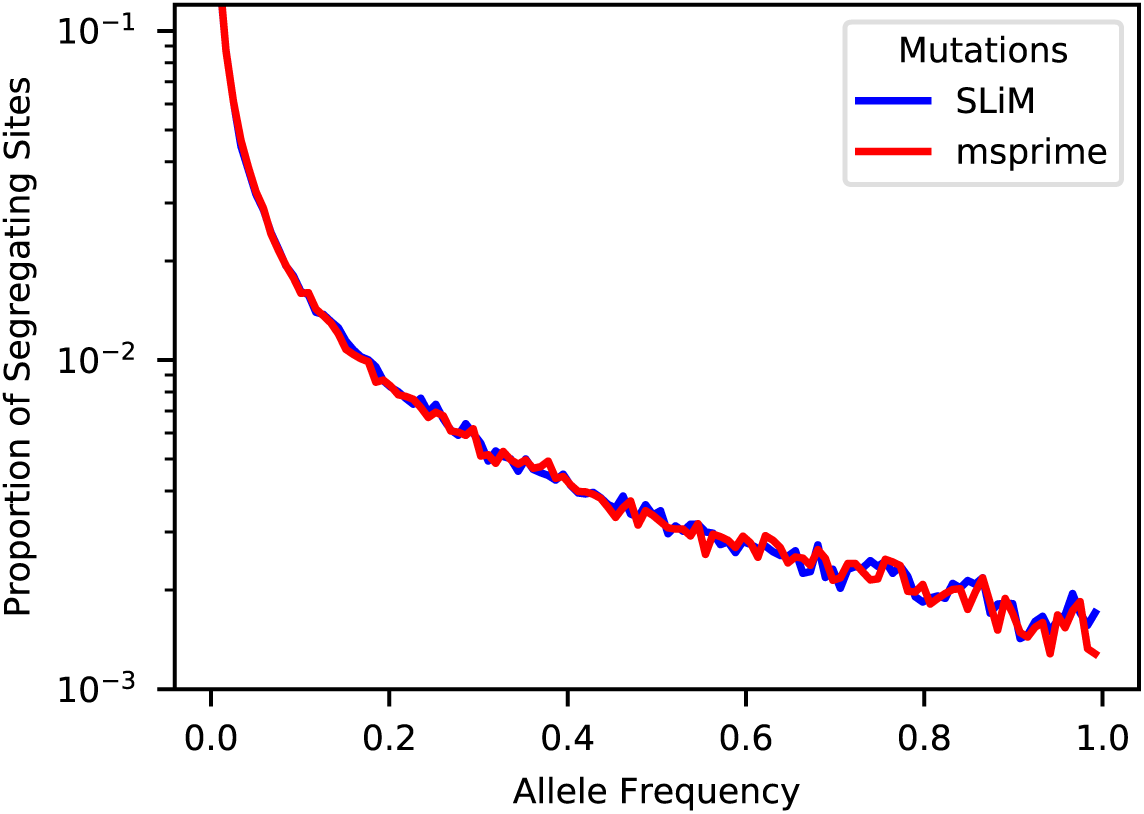
Site frequency spectra from a simulation with neighborhood size = 12.5 when mutations are recorded directly in SLiM (blue line) or applied later in msprime (red line).

**Figure S4.**
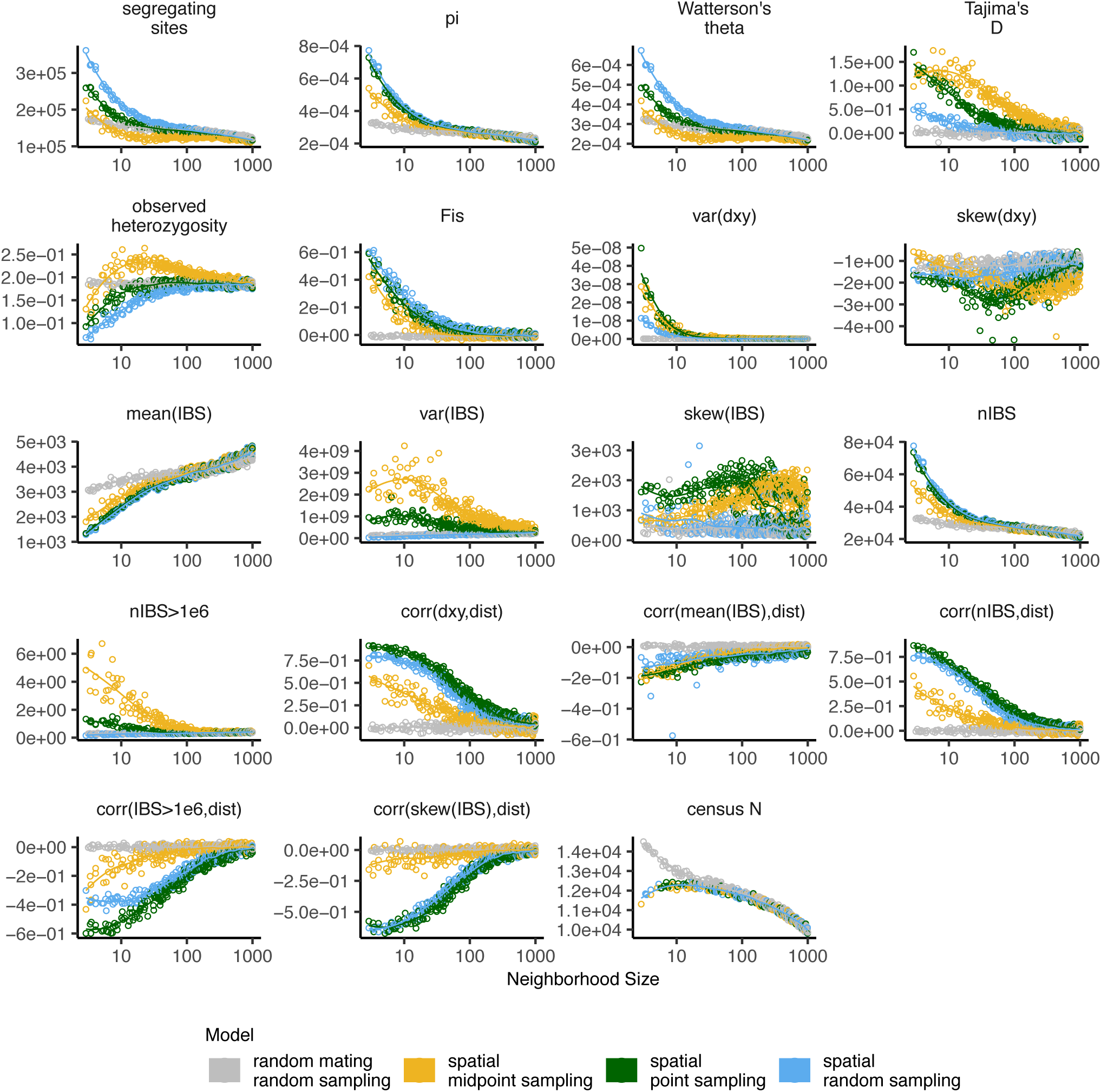
Change in summary statistics by neighborhood size and sampling scheme calculated from simulated sequence data of 60 individuals.

**Figure S5.**
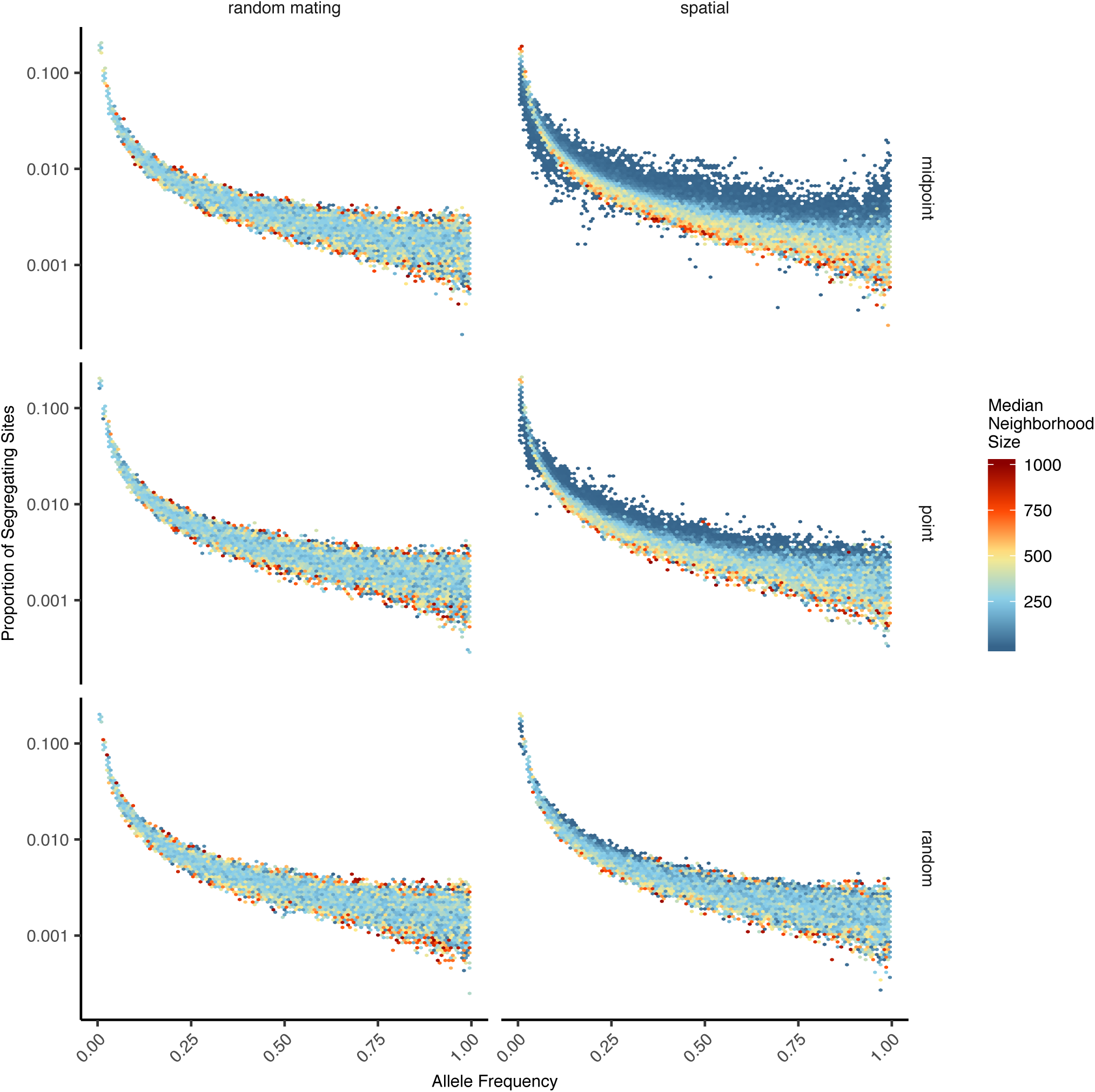
Site frequency spectra for random mating and spatial SLiM models under all sampling schemes.

**Figure S6.**
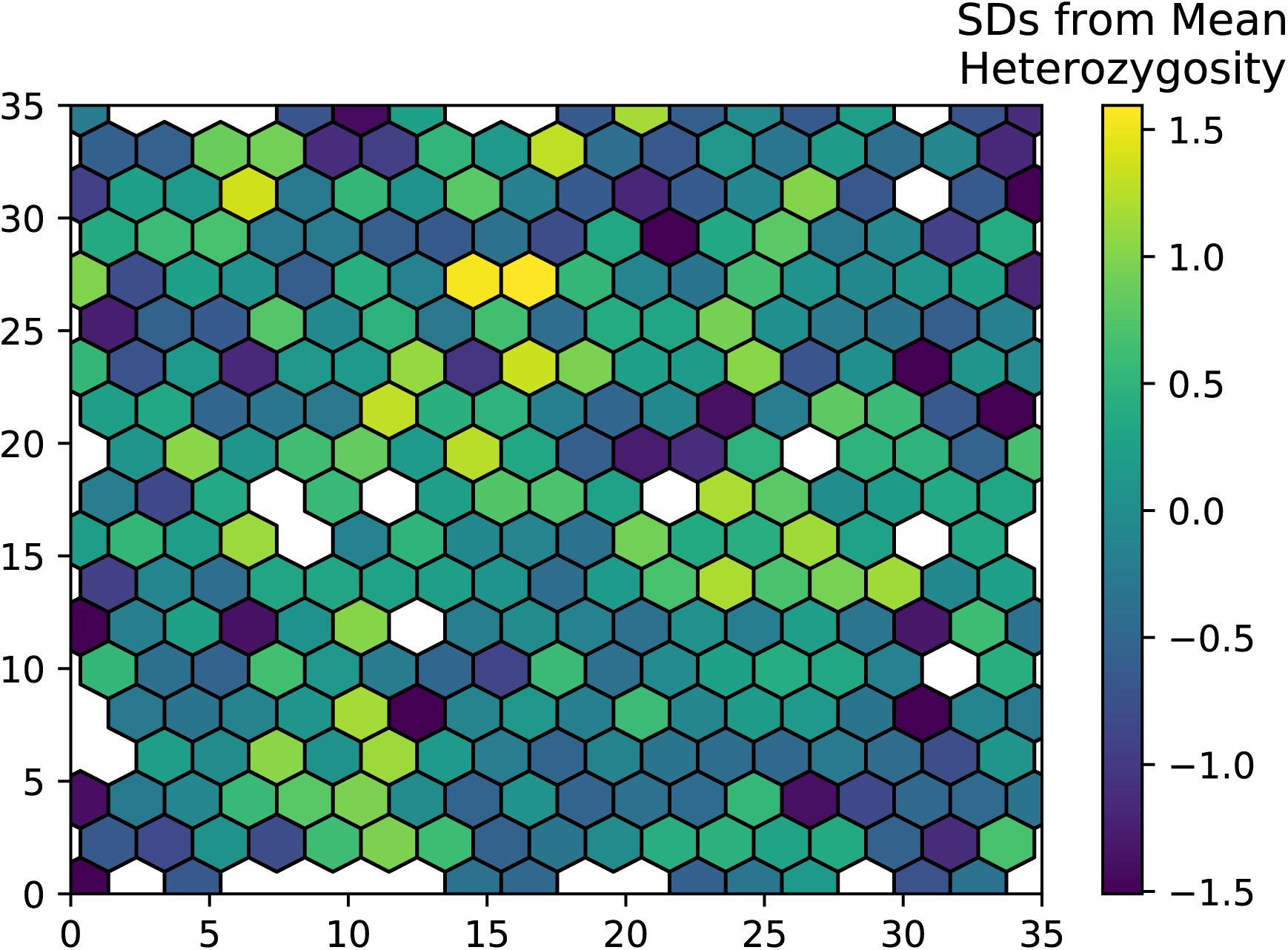
Variation in observed heterozygosity (i.e. proportion of heterozygous individuals) in hexagonal bins across the landscape, estimated from a random sample of 200 individuals from the final generation of a simulation with neighborhood size ≈25. Values were Z-normalized for plotting.

**Figure S7.**
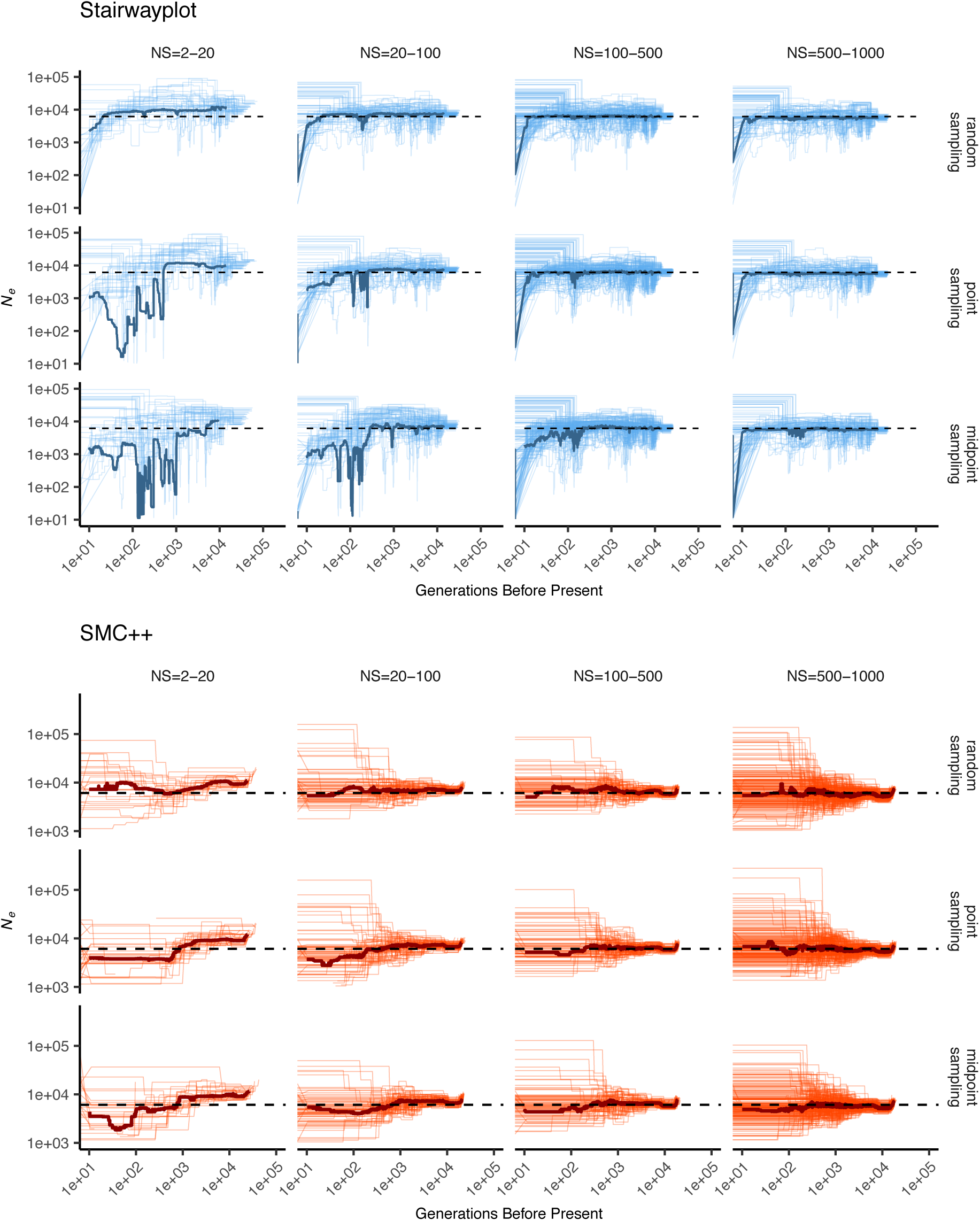
Inferred demographic histories for spatial SLiM simulations, by sampling scheme and neighborhood size (NS) range. Thick lines are rolling medians across all simulations in a bin and thin lines are best fit models for each simulation. Dashed horizontal lines are the average *N*_*e*_ across random-mating SLiM models estimated from *θ*_*π*_.

**Figure S8.**
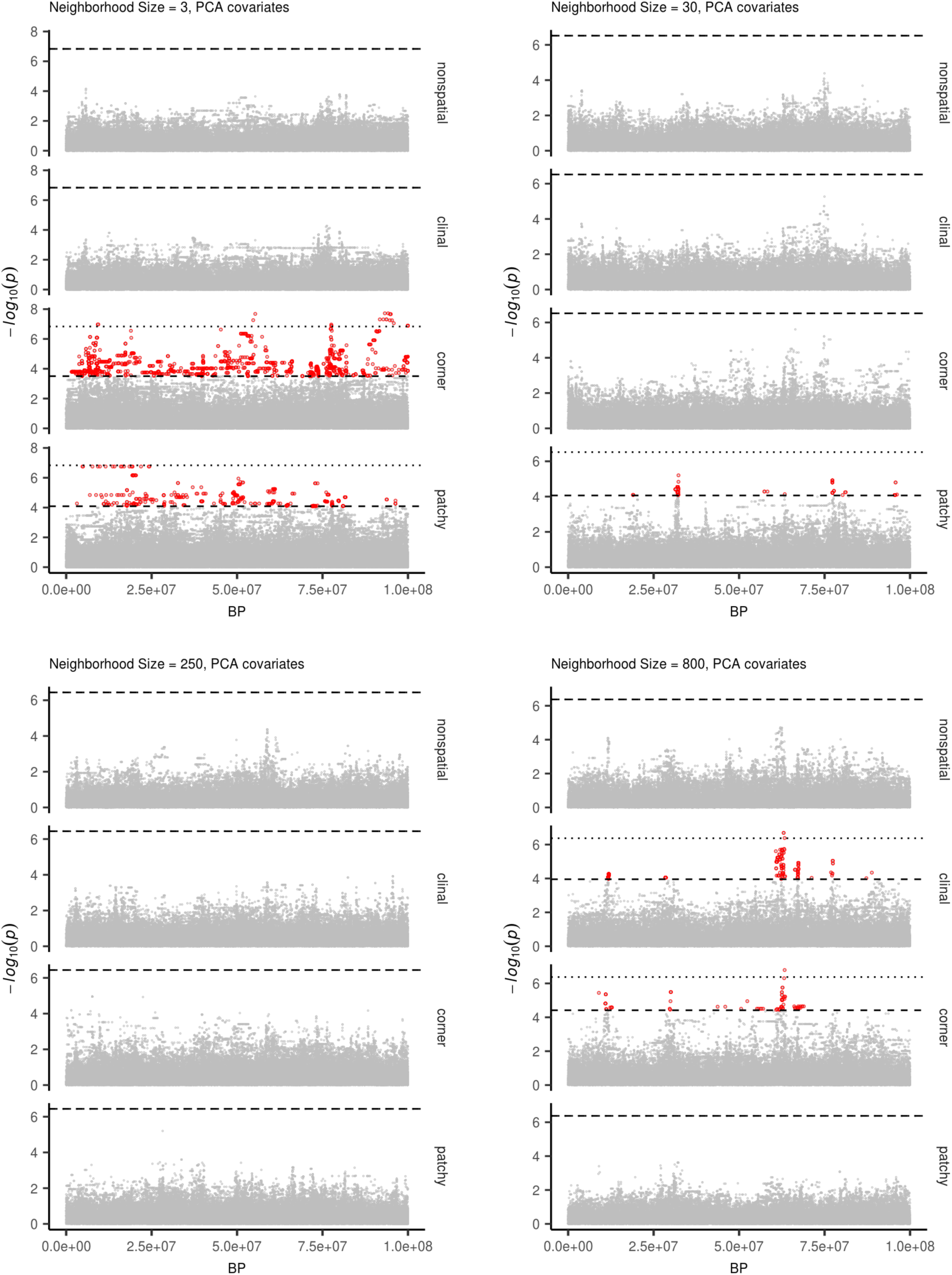
Manhattan plots for a sample of simulations at varying neighborhood sizes. Labels on the right of each plot describes the spatial distribution of environmental factors (described in the methods section of the main text). Points in red are significantly associated with a nongenetic phenotype using a 5% FDR threshold (dashed line). For runs with significant associations the dotted line is a Bonferroni-adjusted cutoff for *p* = 0.05.

**Figure S9.**
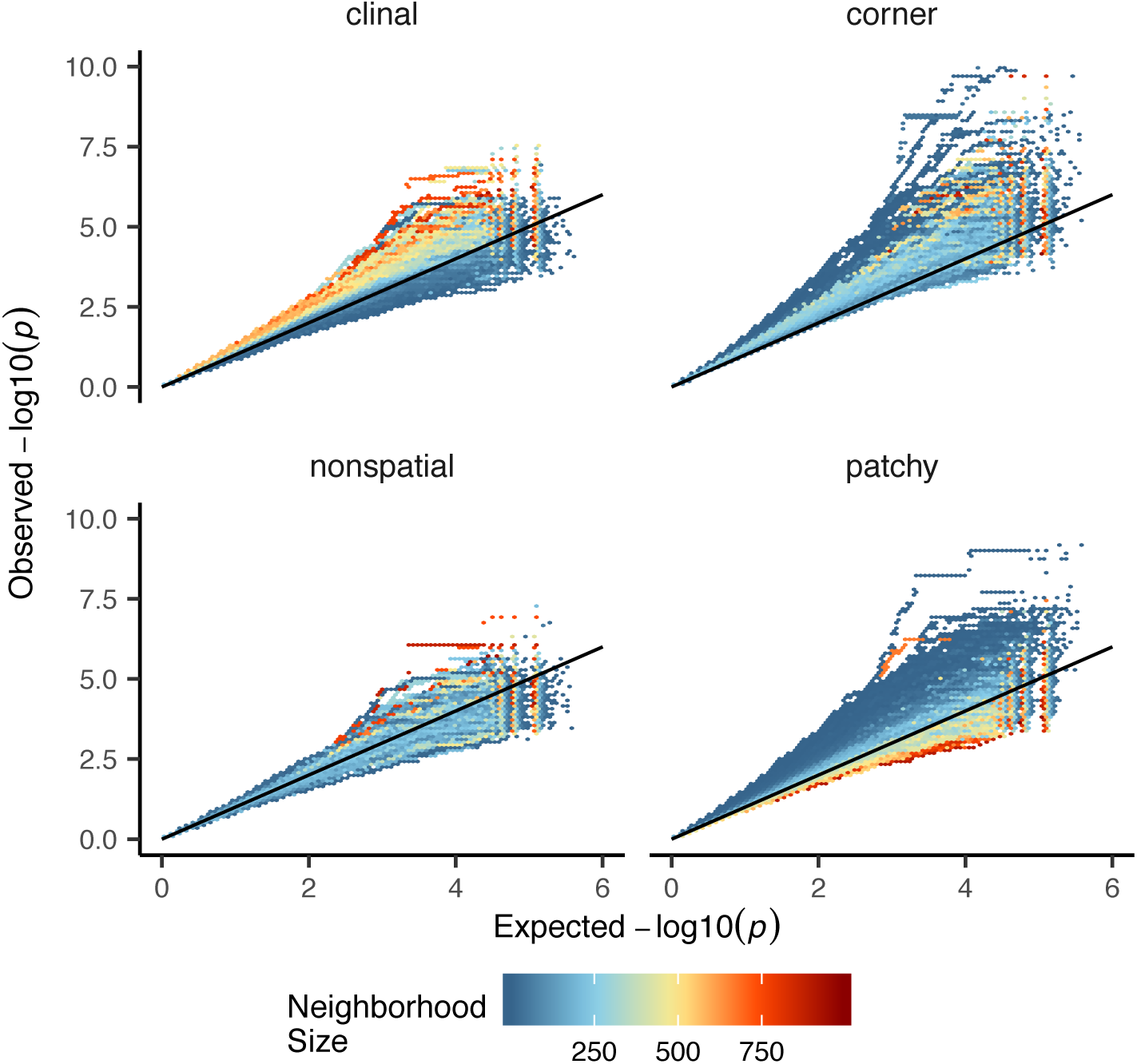
Quantile-quantile plots showing observed and expected − *log*10(*p*) for PC-corrected GWAS run on simulations with varying neighborhood sizes and environmental distributions. Hexagonal bins are colored by the average neighborhood size of simulations with points falling in a given region of quantile-quantile space. Qqplots for a subset of these simulations are shown as lines in Figure 8D.

**Table S1.** Summary statistics calculated on simulated genotypes.

| Statistic | Description |
| --- | --- |
| $\Theta_{/pi}$ | Mean of the distribution of pairwise genetic differences |
| $\Theta_{/W}$ | Effective population size based on segregating sites |
| Segregating Sites | Total number of segregating sites in the sample |
| Tajima's $D$ | Difference in $\Theta_{/pi}$ and $\Theta_{/W}$ over its standard deviation |
| Observed Heterozygosity | Proportion of heterozygous individuals in the sample |
| $F_{IS}$ | Wright's inbreeding coefficient $1 - H_e / H_o$ |
| $var(D_{xy})$ | Variance in the distribution of pairwise genetic distances |
| $skew(D_{xy})$ | Skew of the distribution of pairwise genetic distances |
| $mean(IBS)$ | Mean of the distribution of pairwise identical-by-state (IBS) tract lengths taken over all pairs. |
| $var(IBS)$ | Variance of the distribution of pairwise identical-by-state (IBS) tract lengths taken over all pairs. |
| $skew(IBS)$ | Skew of the distribution of pairwise identical-by-state (IBS) tract lengths taken over all pairs. |
| $nIBS$ | Mean number of IBS tracts with length $> 2bp$ across all pairs in the sample. |
| $nIBS > 1e6$ | Mean number of IBS tracts over $1 \times 10^6bp$ per pair across all pairs in the sample. |
| $corr(D_{xy}, dist)$ | Pearson correlation between genetic distance and $\log_{10}(spatial\ distance)$ |
| $corr(mean(IBS), dist)$ | Pearson correlation between the mean of the IBS tract distribution for each pair of samples and $\log_{10}(spatial\ distance)$ |
| $corr(nIBS, dist)$ | Pearson correlation between the number of IBS tracts for each pair of samples and $\log_{10}(spatial\ distance)$ |
| $corr(IBS > 1e6, dist)$ | Pearson correlation between the number of IBS tracts $> 1 \times 10^6bp$ for each pair of samples and $\log_{10}(spatial\ distance)$ |
| $corr(skew(IBS), dist)$ | Pearson correlation between the skew of the distribution of pairwise haplotype block lengths for each pair of samples and $\log_{10}(spatial\ distance)$ |

**Table S2.** Anova and Levene’s test *p* values for differences by sampling strategy. Bolded values are rejected at *α* = 0.05.

| variable | model | p(equal means) | p(equal variance) |
| --- | --- | --- | --- |
| segsites | random mating | 0.998190 | 0.980730 |
| $\Theta_\pi$ | random mating | 0.997750 | 0.996450 |
| $\Theta_W$ | random mating | 0.998190 | 0.980730 |
| Tajima's $D$ | random mating | 0.879690 | 0.188770 |
| observed heterozygosity | random mating | 0.531540 | 0.433230 |
| $F_{IS}$ | random mating | 0.474790 | 0.785730 |
| $mean(D_{xy})$ | random mating | 0.997770 | 0.996510 |
| $var(D_{xy})$ | random mating | 0.283630 | 0.647240 |
| $skew(D_{xy})$ | random mating | 0.958320 | 0.260750 |
| $corr(D_{xy}, dist)$ | random mating | 0.601980 | **0.000000** |
| $mean(IBS)$ | random mating | 0.997960 | 0.997730 |
| $var(IBS)$ | random mating | 0.486450 | 0.399490 |
| $skew(IBS)$ | random mating | 0.117980 | 0.069770 |
| $nIBS$ | random mating | 0.997680 | 0.996570 |
| $nIBS > 1e6$ | random mating | 0.834870 | 0.888730 |
| $corr(mean(IBS), dist)$ | random mating | 0.073270 | 0.308420 |
| $corr(IBS > 1e6, dist)$ | random mating | 0.268440 | **0.002100** |
| $corr(skew(IBS), dist)$ | random mating | 0.396920 | **0.000620** |
| $corr(nIBS, dist)$ | random mating | 0.581090 | **0.000000** |
| segsites | spatial | **0.000000** | **0.000000** |
| $\Theta_\pi$ | spatial | **0.026510** | **0.013440** |
| $\Theta_W$ | spatial | **0.000000** | **0.000000** |
| Tajima's $D$ | spatial | **0.000000** | **0.000000** |
| observed heterozygosity | spatial | **0.000000** | **0.000000** |
| $F_{IS}$ | spatial | **0.000000** | **0.000120** |
| $mean(D_{xy})$ | spatial | **0.025390** | **0.012910** |
| $var(D_{xy})$ | spatial | **0.004970** | **0.006230** |
| $skew(D_{xy})$ | spatial | **0.000000** | **0.000000** |
| $corr(D_{xy}, dist)$ | spatial | **0.000000** | **0.000000** |
| $mean(IBS)$ | spatial | 0.272400 | 0.114250 |
| $var(IBS)$ | spatial | **0.000000** | **0.000000** |
| $skew(IBS)$ | spatial | **0.000000** | **0.000000** |
| $nIBS$ | spatial | **0.033920** | **0.016640** |
| $nIBS > 1e6$ | spatial | **0.000000** | **0.000000** |
| $corr(mean(IBS), dist)$ | spatial | **0.000000** | 0.590540 |
| $corr(IBS > 1e6, dist)$ | spatial | **0.000000** | **0.000000** |
| $corr(skew(IBS), dist)$ | spatial | **0.000000** | **0.000000** |
| $corr(nIBS, dist)$ | spatial | **0.000000** | **0.000000** |

## Notes

#### Summary of Updates

The main changes we have made have to do with our choice of model. We have added some more discussion of the biological motivation for the particular model of continuous space we use (with references to the ecological literature, where it appears) to the Methods (around (p. 3, l. 131)), and in the Discussion (at (p. 21, l. 744)). We have included more explanation of some of the more opaque consequences of a more realistic demographic model, around (p. 8, l. 343). Finally, we have added a substantial comparison to the discrete stepping-stone model, most of which is in the Appendix.

